# Differential plasmacytoid dendritic cell phenotype and type I Interferon response in asymptomatic and severe COVID-19 infection

**DOI:** 10.1101/2021.04.17.440278

**Authors:** Martina Severa, Roberta A. Diotti, Marilena P. Etna, Fabiana Rizzo, Stefano Fiore, Daniela Ricci, Marco Iannetta, Alessandro Sinigaglia, Alessandra Lodi, Nicasio Mancini, Elena Criscuolo, Massimo Clementi, Massimo Andreoni, Stefano Balducci, Luisa Barzon, Paola Stefanelli, Nicola Clementi, Eliana M. Coccia

## Abstract

SARS-CoV-2 fine-tunes the interferon (IFN)-induced antiviral responses, which play a key role in preventing coronavirus disease 2019 (COVID-19) progression. Indeed, critically ill patients show an impaired type I IFN response accompanied by elevated inflammatory cytokine and chemokine levels, responsible for cell and tissue damage and associated multi-organ failure.

Here, the early interaction between SARS-CoV-2 and immune cells was investigated by interrogating an *in vitro* human peripheral blood mononuclear cell (PBMC)-based experimental model. We found that, even in absence of a productive viral replication, the virus mediates a vigorous TLR7/8-dependent production of both type I and III IFNs and inflammatory cytokines and chemokines, known to contribute to the cytokine storm observed in COVID-19. Interestingly, we observed how virus-induced type I IFN secreted by PBMC enhances anti-viral response in infected lung epithelial cells, thus, inhibiting viral replication. This type I IFN was released by plasmacytoid dendritic cells (pDC) *via* an ACE-2-indipendent mechanism. Viral sensing regulates pDC phenotype by inducing cell surface expression of PD-L1 marker, a feature of type I IFN producing cells. Coherently to what observed *in vitro*, asymptomatic SARS-CoV-2 infected subjects displayed a similar pDC phenotype associated to a very high serum type I IFN level and induction of anti-viral IFN-stimulated genes in PBMC. Conversely, hospitalized patients with severe COVID-19 display very low frequency of circulating pDC with an inflammatory phenotype and high levels of chemokines and pro-inflammatory cytokines in serum.

This study further shed light on the early events resulting from the interaction between SARS-CoV-2 and immune cells occurring *in vitro* and confirmed *ex vivo*. These observations can improve our understanding on the contribution of pDC/type I IFN axis in the regulation of the anti-viral state in asymptomatic and severe COVID-19 patients.

**Author summary:** SARS-CoV-2 pandemic has resulted in millions of infections and deaths worldwide, yet the role of host innate immune responses in COVID-19 pathogenesis remains only partially characterized. Innate immunity represents the first line of host defense against viruses. Upon viral recognition, the secretion of type I and III interferons (IFN) establishes the cellular state of viral resistance, and contributes to induce the specific adaptive immune responses. Moving from *in vitro* evidences on the protective role played by plasmacytoid dendritic cells (pDC)-released type I IFN in the early phase of SARS-CoV-2 infection, here we characterized *ex vivo* the pDC phenotype and the balance between anti-viral and pro-inflammatory cytokines of COVID-19 patients stratified according to disease severity. Our study confirms in COVID-19 the crucial and protective role of pDC/type I IFN axis, whose deeper understanding may contribute to the development of novel pharmacological strategies and/or host-directed therapies aimed at boosting pDC response since the early phases of SARS-CoV-2 infection.

## INTRODUCTION

As of end of March, 2021, COVID-19 pandemic, caused by severe acute respiratory syndrome coronavirus 2 (SARS-CoV-2), has resulted in more than one hundred and twenty million cases and more than two million and six hundred deaths globally (https://www.who.int/publications/m/item/weekly-epidemiological-update-on-covid-19---31march-2021).

The vast majority of SARS-CoV-2 infected subjects are asymptomatic or experience mild to moderate disease, whereas 5 to 10% of them progress, unfortunately, to severe or critical disease, requiring mechanical ventilation and admission to intensive care unit (1, 2). Host characteristics, including age (immunosenescence) and comorbidities (hypertension, diabetes mellitus, lung and heart diseases) may influence the course of the disease (3). Severe pneumonia caused by SARS-CoV-2 is marked by immune system dysfunction and systemic hyperinflammation leading to acute respiratory distress syndrome, macrophage activation, hypercytokinemia and coagulopathy (4).

Interestingly, SARS-CoV-2 infection can be asymptomatic with a rate of presentation estimated to be at least around 17% of total cases (5). Understanding the immunological features of these subjects is challenging but is of key importance to comprehend early events controlling SARS-CoV-2 replication.

Progression and severity of SARS-CoV-2 infection are, indeed, strictly related to virus-triggered immune dysfunctions often associated to defects in type I Interferon (IFN) production, and to over-production of pro-inflammatory cytokines, strongly implicated in the resulting damage to the airways and to the organs (6). Therefore, disease severity is not only due to the virus direct damage but strongly depends on the host inflammatory response to the infection (6–8).

Notably, coronaviruses have evolved several strategies to inhibit IFN anti-viral action triggering many escaping strategies (9, 10). This feature is relevant for SARS-CoV-2, which replicates more efficiently than SARS-Co-V (11), but induces less release of type I, II and III IFNs (7, 12). Hence, different SARS-CoV-2-encoded proteins can limit type I IFN functions or pathways (13). In particular, nsp13, nsp14, nsp15, ORF6, ORF8, ORF3b and nucleocapsid proteins act as competent suppressors of IFN-β; while nsp1, nsp12, nsp13, nsp15 and the M protein are also potent inhibitors of the MAVS pathway (14–16).

In this paper, we investigate the SARS-CoV-2-elicited innate immune response using a human PBMC-based *in vitro* experimental model. We evaluate the key role of plasmacytoid dendritic cells (pDC) in the induction of type I IFN response in this context. Also, we performed an in-depth analysis of pDC phenotype and IFN and cytokine levels in COVID-19 patients, stratified according to disease severity, to evaluate possible correlation with COVID-19 progression and subsequent life-threatening complications.

## RESULTS

### SARS-CoV-2 stimulation of human PBMC induces a TLR7/8-dependent cytokine and chemokine production *in vitro*

To understand if cells of immune system can sense SARS-CoV-2 leading to activation of innate response, we stimulated human PBMC with a clinical SARS-CoV-2 isolate.

To monitor the virus kinetic after incubation with human PBMC, we first treated these cells with SARS-CoV-2 at different multiplicity of infection (MOI; 0.01, 0.02, 0.04 and 0.1). Viral titration in supernatants from PBMC cultures, performed at the inoculum check at time zero (T0), as well as at 24 (T1) and 48 (T2) hours post viral infection, showed no viral replication (**Fig 1**), in spite of cell viability that was not altered (**S1 Table**). These results indicate that PBMC, which are indeed not natural target cells of SARS-CoV-2, are not permissive, at least *in vitro*, to infection.

**1 Fig.**
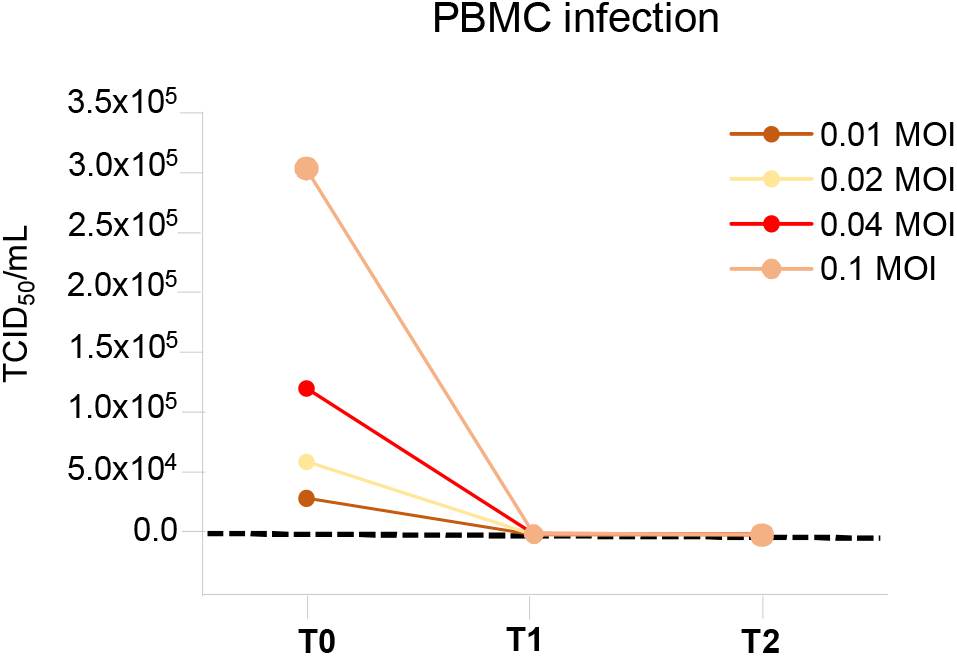
Human PBMC are not permissive to SARS-CoV-2 infection. Virus titers were evaluated in viral inoculum (T0) as well as in peripheral blood mononuclear cell (PBMC) supernatants after 24 (T1) and 48 hours (T2) post SARS-CoV-2 infection in cultures treated at a multiplicity of infection (MOI) of 0.01, 0.02, 0.04 and 0.1. Shown results were calculated by Endpoint Dilution Assay by using the Reed-Muench formula and reported as TCID_50_/ml and derived from three experiments conducted separately.

Having in mind that SARS-CoV-2 infection of both upper and lower airways regulates type I and type III IFN responses (12), here we studied by real-time PCR the expression of these anti-viral cytokines in the mixed immune cell population of PBMC and found a dose-dependent induction of high levels of type I IFN-αs and IFN-β, as well as type III IFN-λ1 transcripts, as compared to cells stimulated with the TLR7/8 ligand R848, used as positive control (**Fig 2A**). In line with these data, we found a strong release of IFN-αs in culture supernatants (**Fig 2B**) and a concomitant induction of Mx dynamin-like GTPase 1 (Mx1) expression (**Fig 2C**), one of the classical IFN-inducible genes representing the so-called anti-viral IFN-signature. Thus, in human PBMC SARS-CoV-2 was unable to interfere efficiently with the intracellular machinery responsible for anti-viral IFN release and induction of IFN signature. Moreover, the addition of a specific inhibitor of TLR7/8 signaling (I-TLR7/8) completely abolished SARS-CoV-2-induced type I IFN response in PBMC (**Fig 2**), suggesting that the viral RNA may be involved in induction of IFN production in this setting. The level of cytokines and gene expression was unchanged in mock-treated cultures (**Fig 2**). Also, in the same experimental setting we found a remarkable production of inflammatory cytokines such as IL-6, TNF-α and IL-1β, which are well-recognized players of the cytokine storm occurring in COVID-19 patients (17), while IL-12p70 level was reduced in SARS-CoV-2-stimulated PBMC (**Fig 3A**). Importantly, the anti-inflammatory IL-10 factor was also induced but its level was inversely related to virus MOI. In particular, IL-10 heavily decreased at higher MOI where inflammatory cytokines were enhanced (**Fig 3A**). Therefore, as for IFNs, the production of pro-inflammatory cytokines is dependent on TLR7/8 signaling (**Fig 3A**).

**2 Fig.**
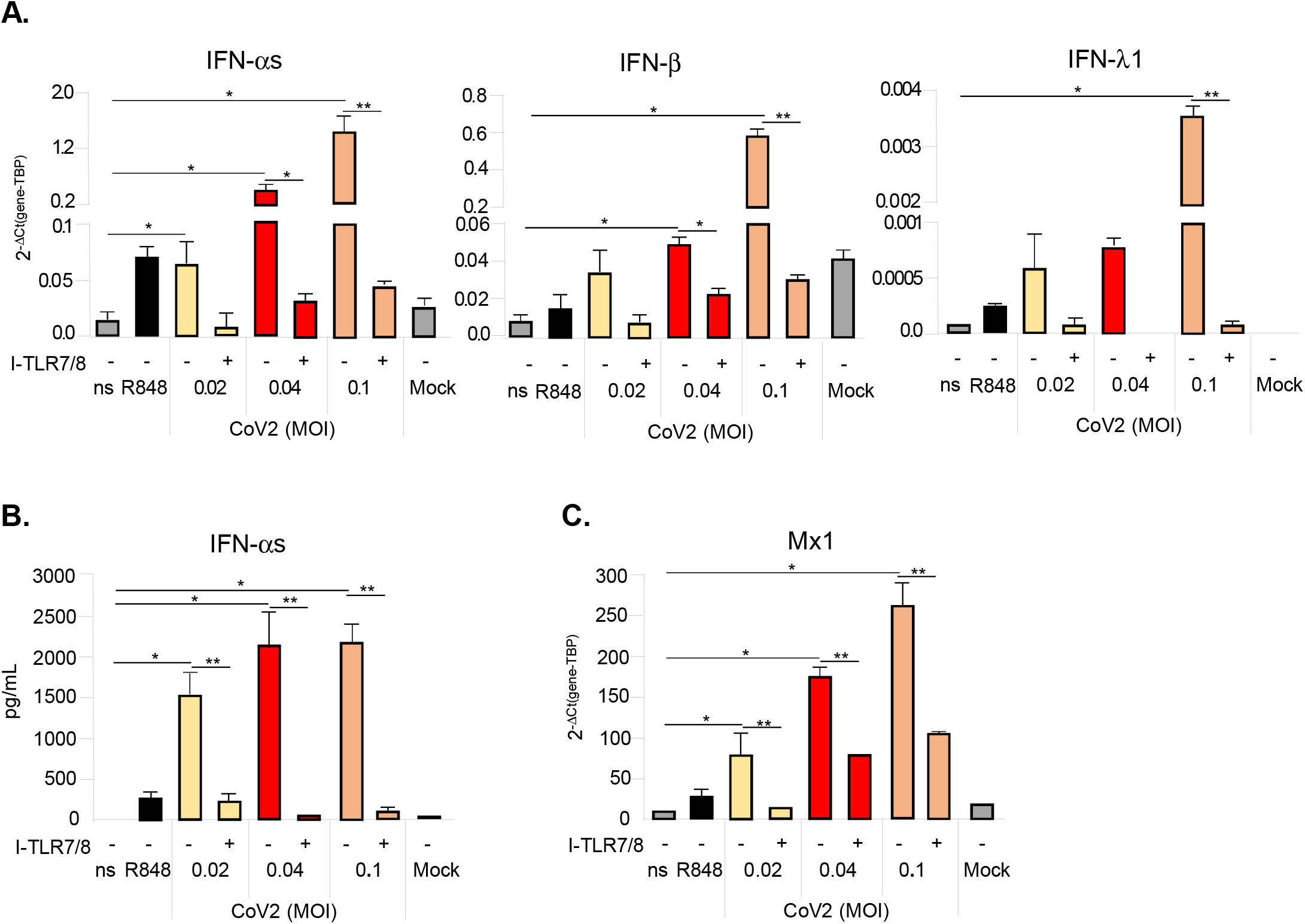
SARS-CoV-2 stimulation induces a TLR7/8-dependent type I and III IFN response in PBMC. Peripheral blood mononuclear cells (PBMC) were left untreated (not stimulated, ns) or stimulated for 24 hours with the TLR7/8 agonist R848 (5μM) as positive control, with SARS-CoV-2 (CoV2) at different multiplicity of infection (MOI; 0.02, 0.04 and 0.1) in presence or absence of a 30 minute pre-treatment with a specific TLR7/8 inhibitor (I-TLR7/8, 1μM) or with Mock medium only. (**A**) Relative expression of IFN-αs, IFN-β, IFN-λ1 genes was measured by quantitative real time PCR analysis. All quantification data are normalized to TBP level by using the equation 2^−ΔCt^. (**B**) Production of IFN-αs was tested by specific ELISA in 24 hour-collected cell culture supernatants. (**C**) Mx1 gene expression was quantified by real time PCR as described above. Shown results were mean relative values ± SEM of 5 independent experiments. P-values were depicted as follows: *p≤0.05; ** p≤0.001.

**Fig. 3.**
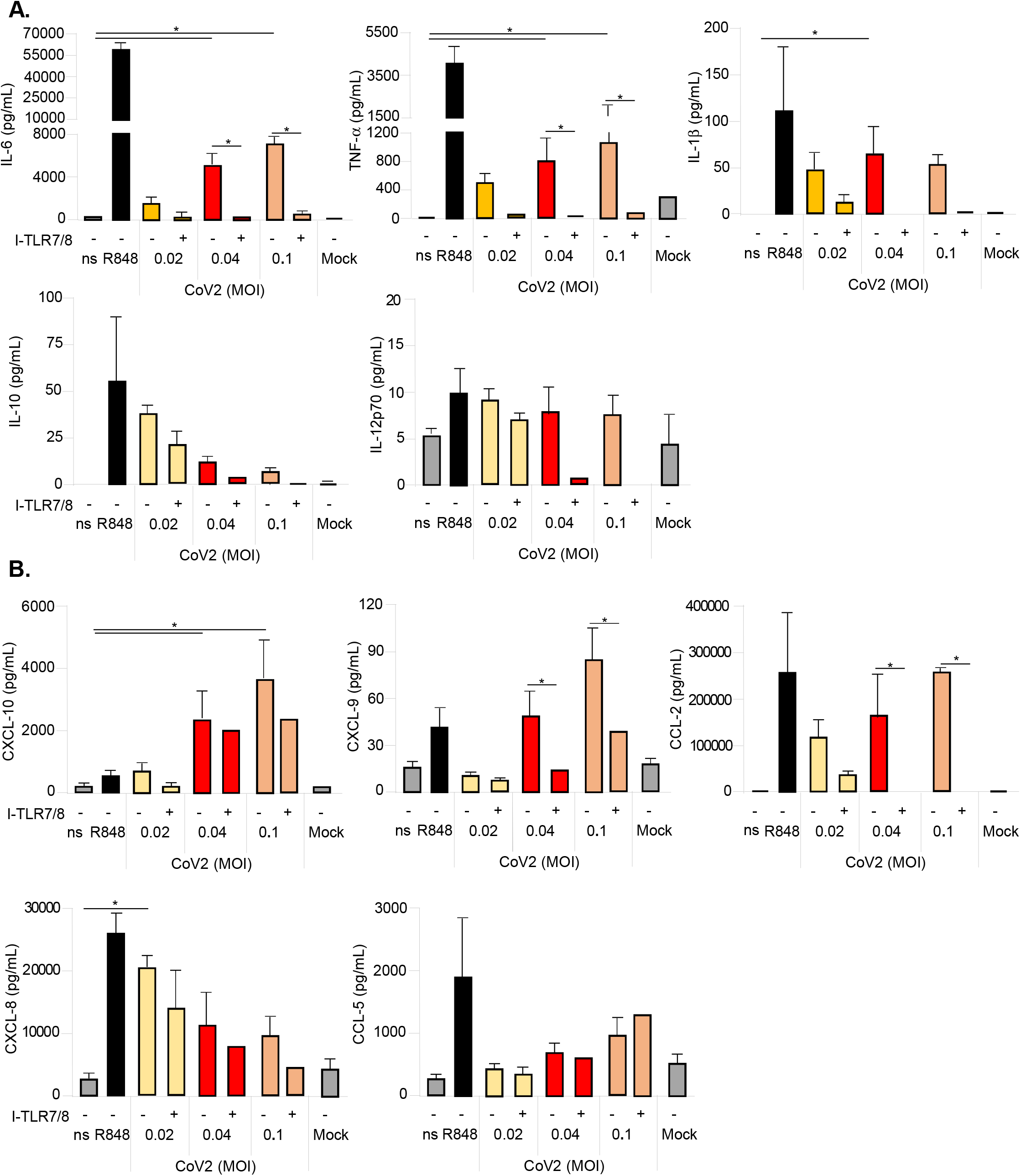
SARS-CoV-2 stimulation induces a TLR7/8-dependent cytokine and chemokine production in PBMC. Peripheral blood mononuclear cells (PBMC) were left untreated (not stimulated, ns) or stimulated for 24 hours with the TLR7/8 agonist R848 (5μM) as positive control, with SARS-CoV-2 (CoV2) at different multiplicity of infection (MOI; 0.02, 0.04 and 0.1) in presence or absence of a 30 minute pre-treatment with a specific TLR7/8 inhibitor (I-TLR7/8, 1μM), or with Mock medium only. Production of cytokines (IL-6, TNF-α, IL-1β, IL-10, IL-12p70) (**A**) and chemokines (CXCL-10, CXCL-9, CCL-2, CXCL-8, CCL-5) (**B**) was tested by multiparametric cytokine bead arrays in collected cell culture supernatants. Shown results were mean relative values ± SEM of 5 independent experiments. P-values were depicted as follows: *p≤0.05.

The recruitment and coordination of specific subsets of leukocytes at the site of viral infection heavily relies on chemokine secretion. Thus, in supernatants of PBMC stimulated with increasing doses of SARS-CoV-2 we also tested the release of CXC motif chemokine ligand 10 (CXCL-10 or IFN-γ inducible protein, IP-10), CXCL-9 (or monokine induced by IFN-γ, MIG), C-C motif chemokine ligand 2 (CCL-2 or macrophage cationic peptide 1, MCP-1), CXCL-8 (or IL-8) and CCL-5 (or Rantes) (**Fig 3B**). Among the studied chemokines, CXCL-10, CXCL-9 and CCL-2 showed the highest induction in a viral dose-dependent manner (**Fig 3B**). In addition, our data showed that CXCL-9 and CCL-2 production was, in particular, TLR7/8-dependent (**Fig 3B**). The release of analyzed cytokines and chemokines was further increased at 48 hours post infection indicating an incremental SARS-CoV-2-dependent stimulation in our PBMC-based culture setting at the tested MOI (0.04 and 0.1) (**S1A and S1B Fig**).

Most importantly, by comparing infectious and UV-inactivated SARS-CoV-2 we also demonstrated that the production of type I IFN and of the pro-inflammatory cytokines IL-6, TNF-α and IL-1β (**Fig 4A**) as well as of the chemotactic factors CXCL-10, CXCL-9 and CCL-2 (**Fig 4B**) required infectious virus.

**4 Fig.**
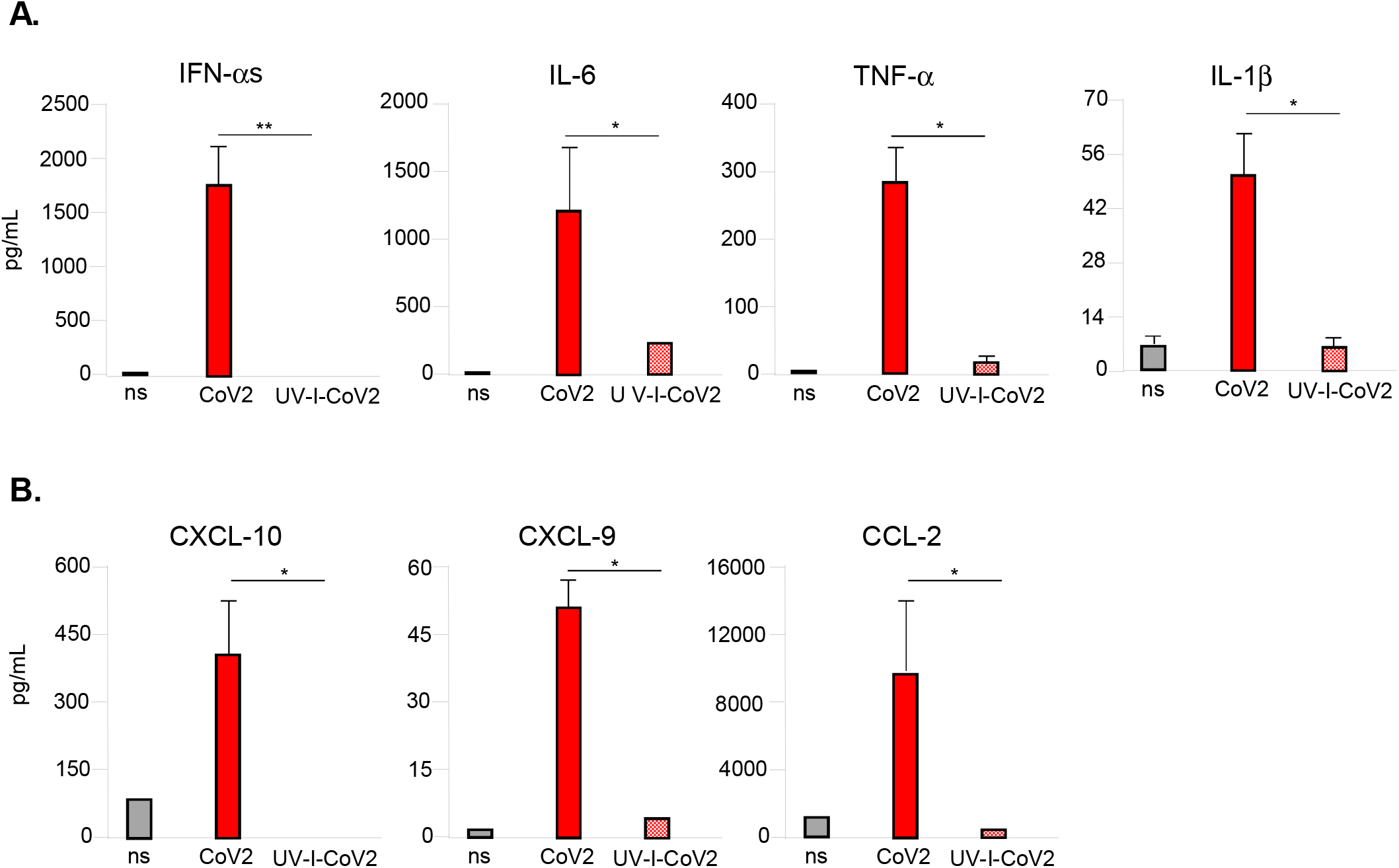
SARS-CoV-2-mediated cytokine and chemokine production is not induced upon PBMC stimulation with UV-inactivated virus. Peripheral blood mononuclear cells (PBMC) were left untreated (not stimulated, ns) or stimulated for 24 hours with infectious or UV-inactivated (UV-I) SARS-CoV-2 (CoV2, MOI=0.04) in presence or absence of a 30 minute pre-treatment with a specific TLR7/8 inhibitor (I-TLR7/8, 1μM). Production of cytokines (IFN-αs, IL-6, TNF-α, IL-1β) and chemokines (CXCL-10, CXCL-9, CCL-2) (**B**) was tested by specific ELISA (IFN-αs only) or multiparametric cytokine bead arrays in collected cell culture supernatants. Shown results were mean relative values ± SEM of 3 independent experiments. P-values were depicted as follows: *p≤0.05.

### SARS-CoV-2-induced type I IFN secreted by human PBMC inhibits virus infection of Calu-3 cells

Several studies have clearly shown that SARS-CoV-2 infection modulates the release of type I and III IFNs (7, 12, 18). In our study we observed that SARS-CoV-2 infection of the lung carcinoma epithelial cell line Calu-3 triggers IL-6 expression while was unable to induce Mx1 gene expression, mirroring a strong reduction of type I IFN production and, thus, confirming what already observed (7, 12) (**Fig 5A**).

**5 Fig.**
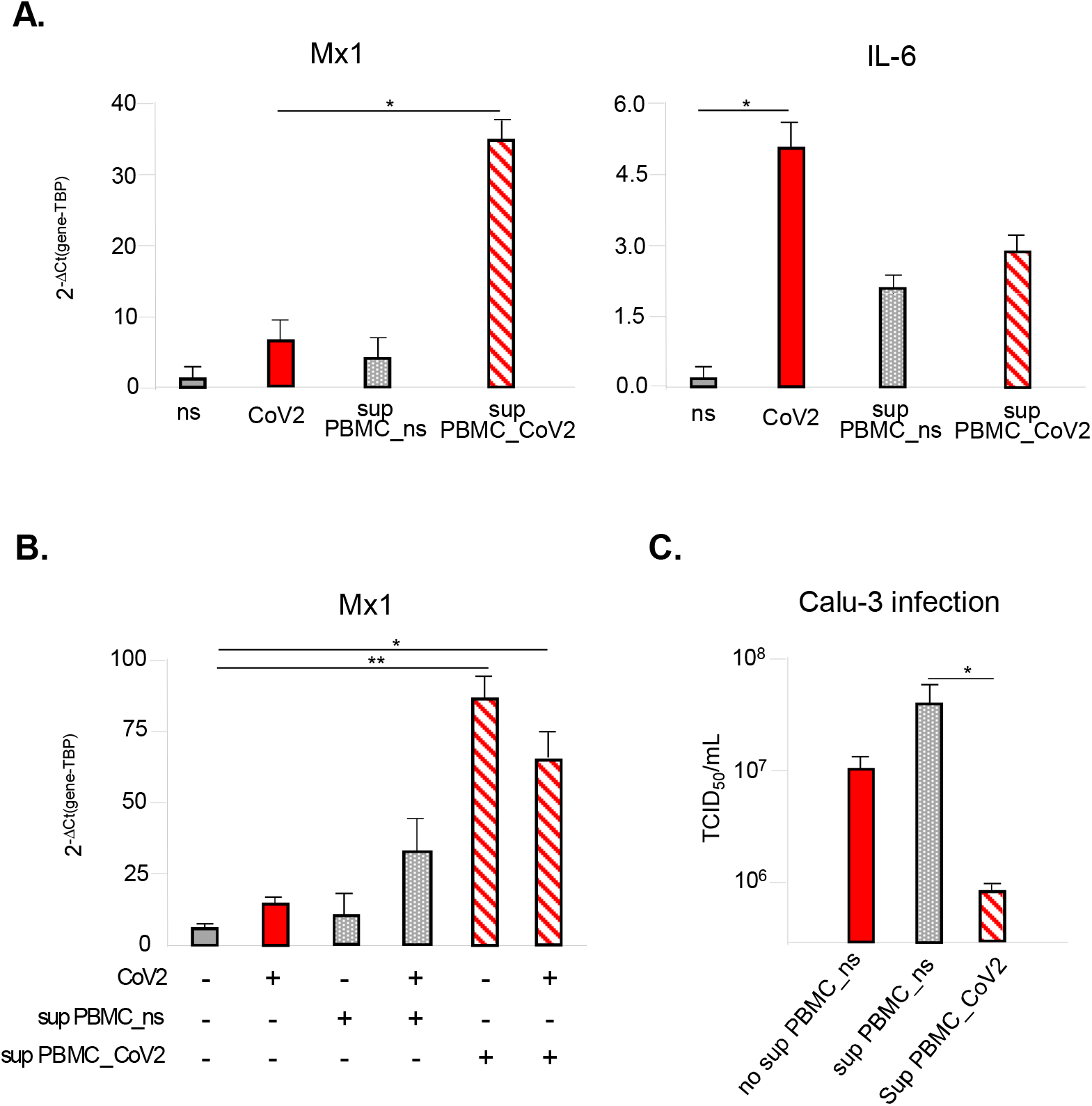
Type I IFN-induced anti-viral state impacts on SARS-CoV-2 infected Calu-3 lung epithelial cell line. (**A**) The human lung epithelial cell line Calu-3 was left untreated (not stimulated, ns), infected for 24 hours with SARS-CoV-2 at the back titration dose calculated in peripheral blood mononuclear cells (PBMC) (10^4^ TCID_50_/ml for 0.02 MOI) or treated with 24 hour-supernatants collected from either ns PBMC cultures (sup PBMC_ns, grey dotted bars) or from SARS-CoV-2 (MOI=0.02)-treated PBMC cultures (sup PBMC_CoV2, red transverse striped bars). Relative expression of Mx1 and IL-6 genes was measured by quantitative real time PCR analysis. All quantification data are normalized to TBP level by using the equation 2^−ΔCt^. (**B**) Calu-3 cell line was infected with SARS-CoV-2 at the back titrated dose calculated in PBMC (10^4^ TCID_50_/ml for 0.02 MOI) in presence of 24 hour-supernatants collected from either ns PBMC cultures (sup PBMC_ns, grey dotted bars) or from SARS-CoV-2 (MOI=0.02)-treated PBMC cultures (sup PBMC_CoV2, red striped bars). Mx1 gene expression was quantified by real time PCR and quantification data normalized to TBP level by using the equation 2^−ΔCt^. (**C**) Virus titers were evaluated in supernatants from infected Calu-3 cultures described in (**B**) by Endpoint Dilution Assay by using the Reed-Muench formula and reported as TCID_50_/ml. Shown results were mean relative values ± SEM of 3 independent experiments. P-values were depicted as follows: *p≤0.05; ** p≤0.001.

Importantly, we first demonstrated that treatment of Calu-3 cells with supernatants collected from SARS-CoV-2-stimulated human PBMC (sup PBMC_CoV2) strongly induced the transcription of the IFN-inducible gene Mx1, differently to what observed in cultures treated with supernatant from not stimulated PBMC (sup PBMC_ns) (**Fig 5A**).

To understand if type I IFN naturally released by infected PBMC can inhibit SARS-CoV-2 replication, we infected Calu-3 cells conditioned with either sup PBMC_ns or sup PBMC_CoV2 (**Fig 5B and 5C**). In our experimental setting, while addition of sup PBMC_ns affected neither Mx1 transcription nor SARS-CoV-2 replication, presence of sup PBMC_CoV2 correlated to an enhanced ISG signature and combined inhibition of SARS-CoV-2 infection as revealed by viral titrations of supernatants from Calu-3 cells (**Fig 5B and 5C**).

Hence, PBMC-derived type I IFN may rescue SARS-CoV2-mediated block of anti-viral response, thus, limiting viral expansion.

### Human pDC produce high levels of type I IFN upon SARS-CoV-2 stimulation

pDC are classified as the major type I IFN producing cells following viral infections by sensing viral RNAs *via* TLR7 (19). Given the TLR7/8-dependent release of the anti-viral cytokines detected in SARS-CoV-2-stimulated PBMC, we hypothesized that pDC could be the main cell type responsible for type I IFN production. In addition, we also tested the possibility that the TLR8-expressing resting CD14^+^ monocytes would be involved.

Thus, pDC were purified from PBMC of healthy donors and stimulated for 24 hours with SARS-CoV-2 (0.1 MOI) (**Fig 6**). In pDC cultures a strong production of IFN-αs, comparable to what found in PBMC, was detected, and this was clearly dependent on TLR7 signaling since I-TLR7/8 blocked release of this cytokine (**Fig 6A**). SARS-CoV-2 stimulation of pDC did not depend on ACE-2 mediated viral entry or on the transmembrane serine protease 2 (TMPRSS2), both not expressed by these cells in baseline condition or after stimulation with IFN-αs and IFN-β (**S2A Fig**). Conversely, this instead occurs in human airway epithelial cells (20) and in Calu-3 lung epithelial cell line (**S2A Fig**).

**6 Fig.**
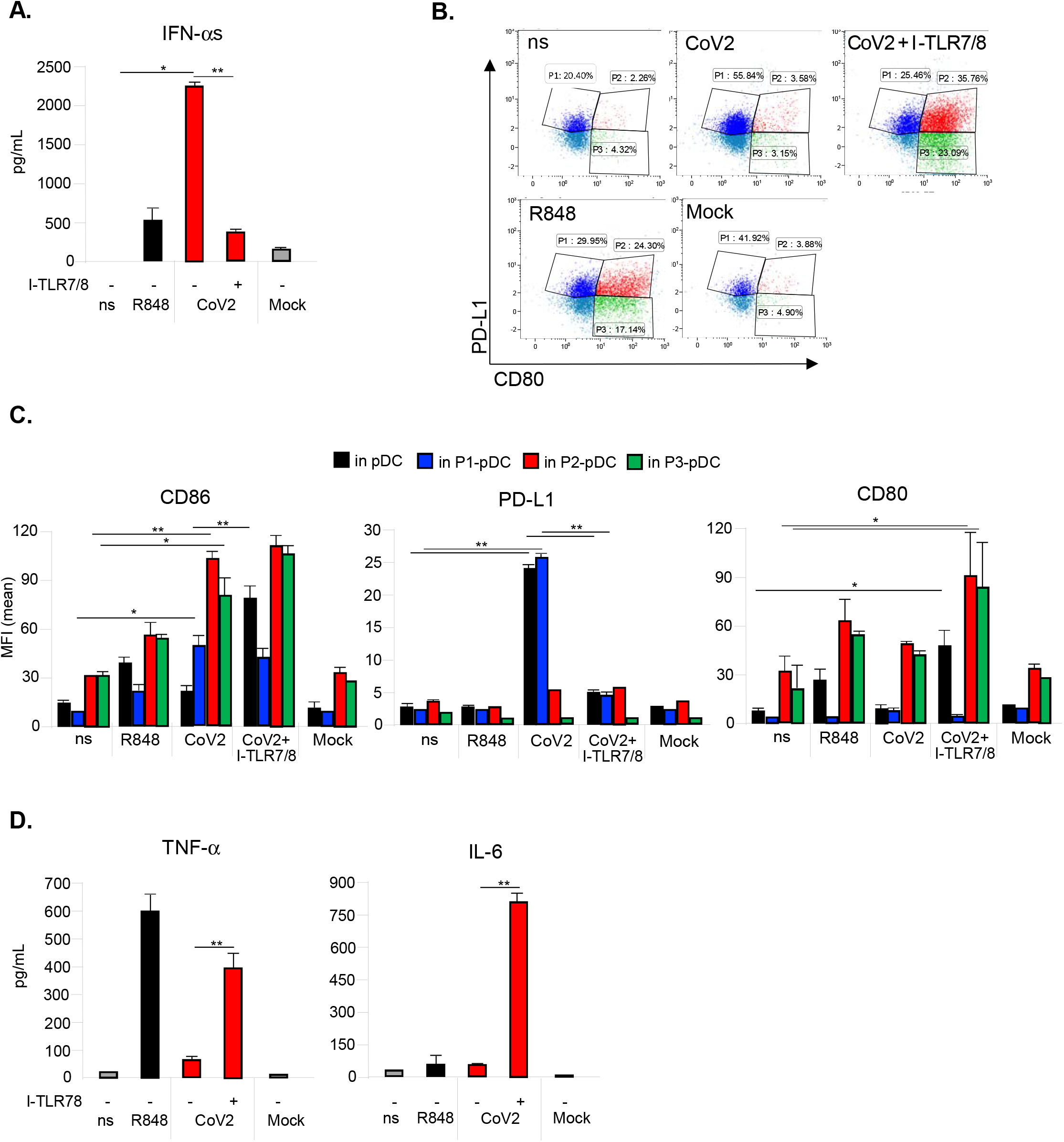
SARS-CoV-2 stimulation drives TLR7-dependent type I IFN production and phenotypical modification in pDC. Purified plasmacytoid dendritic cells (pDC) were left untreated (not stimulated, ns) or stimulated for 24 hours with the TLR7/8 agonist R848 (5μM) as positive control, with SARS-CoV-2 (CoV2; MOI=0.1) in presence or absence of a 30 minute pre-treatment with a specific TLR7/8 inhibitor (I-TLR7/8, 1μM), or with Mock medium only. (**A**) Production of IFN-αs was tested by specific ELISA in 24 hour-collected pDC culture supernatants. pDC stimulated for 24 hours as described above were stained with anti-BDCA4, PD-L1, CD80 and CD86 antibodies. The percentage (%) of pDC sub-populations was evaluated by flow cytometry in live/single BDCA4^+^ pDC; in particular P1-pDC (PD-L1^+^CD80^−^, in blue), P2-pDC (PD-L1^+^CD80^+^, in red) and P3-pDC (PD-L1^−^CD80^+^, in green). Representative dot plot profile out of 3 different experiments independently conducted is shown. (**C**) Surface expression of CD86, PD-L1 and CD80 was determined as mean fluorescence intensity (MFI) by flow cytometer analysis. (**D**) Production of TNF-α and IL-6 was tested by cytometric bead assay in 24 hour-collected pDC culture supernatants. Shown results were mean relative values ± SEM of 3 independent experiments. P-values were depicted as follows: *p≤0.05; ** p≤0.001.

Interestingly, *in vitro* stimulation by SARS-CoV-2 of monocytes, which exclusively express ACE-2 but not TMPRSS2 (**S2A Fig**), did not lead to IFN-α or IL-6 production as compared to R848 treatment (**S2C Fig**), even if these cells can be efficiently infected by the virus (21).

The responsiveness of pDC, monocyte and Calu-3 to exogenous IFN-αs and IFN-β was tested by evaluating the induction of Mx1 to verify the bona fide of ACE-2 and TMPRSS2 expression data (**S2B Fig**).

pDC undergo phenotypical diversification in response to viral infections or single stimuli through environmental plasticity (22). They can diversify into three stable populations: P1-pDC (PD-L1^+^CD80^−^) specialized for type I IFN production, P2-pDC (PD-L1^+^CD80^+^) displaying both innate and adaptive functions and P3-pDC (PD-L1^−^CD80^+^) specifically with adaptive functions (22). In our experimental setting we monitored pDC phenotype and found a clear-cut increase of the frequency of PD-L1^+^ P1-pDC, in accordance with the detected high IFN-α release, upon SARS-CoV-2 stimulation (**Fig 6B**). In line with these data, SARS-CoV-2-treated total pDC and P1-pDC display a very high surface expression of PD-L1 (**Fig 6C**). In presence of the TLR-7/8 inhibitor, P1 phenotype upon SARS-CoV-2 stimulation reverted into the more adaptive P2 and P3 populations, similarly to R848-treated cultures (**Fig 6B**), with a strong increase of the surface level of CD80 (**Fig 6C**). Accordingly, along with the maturation process, exemplified by the induction of the co-stimulatory marker CD86 (**Fig 6C**), SARS-CoV-2-treated pDC turn from type I IFN- into TNF-α-and IL-6-producing cells in the presence of the TLR-7/8 inhibitor (**Fig 6D**).

### pDC of asymptomatic and hospitalized COVID-19 subjects display a different phenotype

Having defined the key role of pDC in the induction of a type I IFN-mediated anti-viral state in SARS-CoV-2-treated human PBMC, we then moved to study this cell type *ex vivo* in PBMC collected from individuals with asymptomatic SARS-CoV-2 infection (CP-AS, n=8), in hospitalized COVID-19 patients (CP, n=6) and in a cohort of healthy donors matched for sex and age (HD, n=5) (see gating strategy in **S3A Fig** and patient characteristics in **S2 and S3 Tables**).

We first monitored whether the degree of COVID-19 severity would match to a differential level of circulating pDC and found a striking difference in the three analyzed groups (**Fig 7A**). The frequency as well as the absolute number of pDC were reduced in asymptomatic SARS-CoV2 infected subjects as compared to HD (**Fig 7A**). According to the observed lymphopenia (**S2 Table**), symptomatic hospitalized patients had a very and consistently low level/depletion of circulating pDC independently of their age and sex (**Fig 7A**).

**7 Fig.**
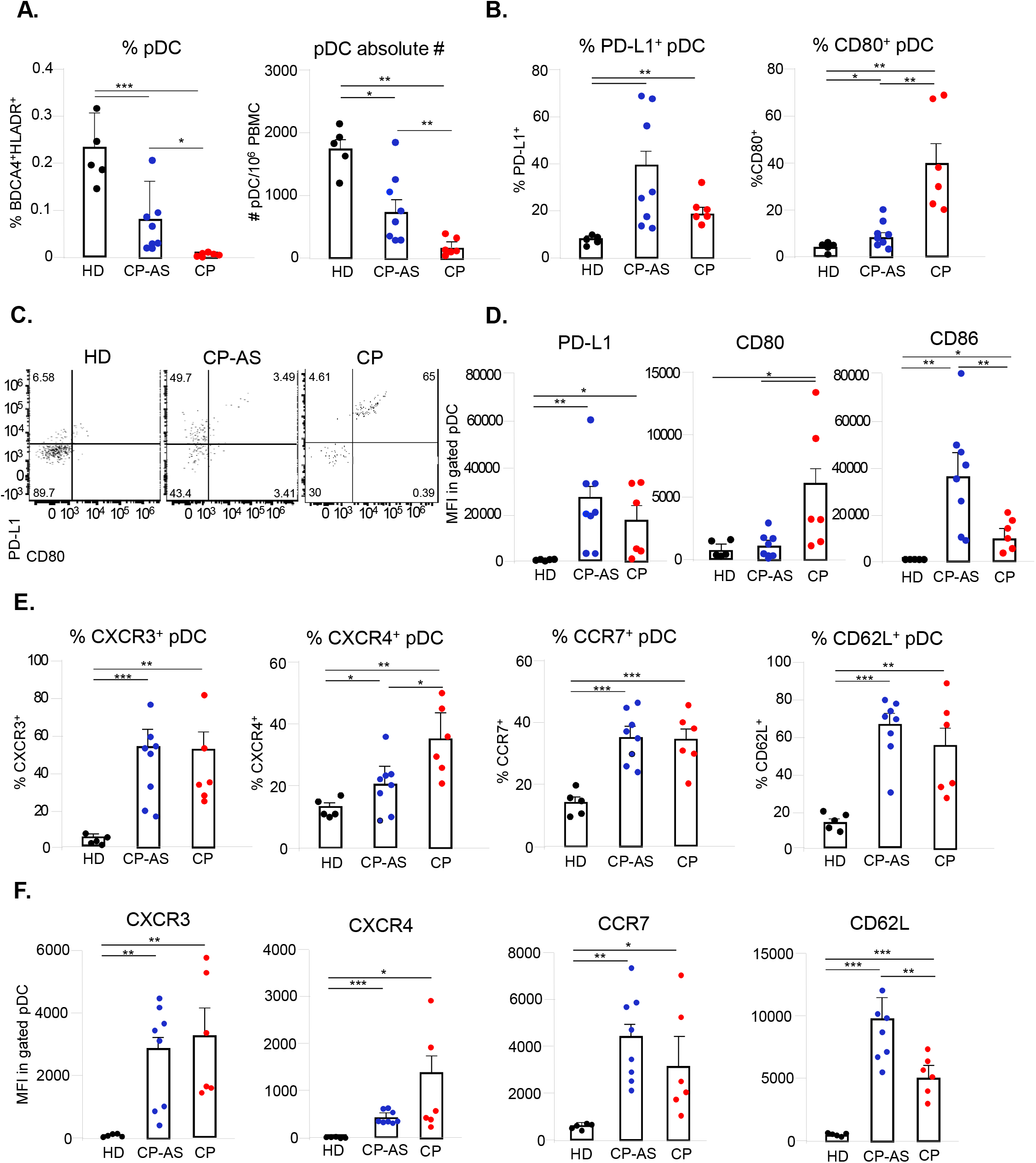
pDC differently activate and express chemokine receptors for homing to SARS-CoV-2 infected tissues in COVID-19 asymptomatic and hospitalized patients. Freshly isolated peripheral blood mononuclear cells (PBMC) from asymptomatic (CP-AS, n=8) and hospitalized COVID-19 patients (CP, n=6) as well as matched healthy donors (HD, n=5) were stained with a cocktails of antibodies to study by flow cytometry plasmacytoid dendritic cell (pDC) frequency and absolute number (**A**), diversification and activation status (Lineage, CD123, BDCA-4, HLA-DR, PD-L1, CD80 and CD86) (**B, C, D**), as well as expression of chemokine receptors (Lineage, CD123, BDCA-4, HLA-DR, CXCR4, CXCR3, CCR7 and CD62-L) (**E, F**). The percentage (%) of shown pDC sub-populations (**B, C, E**) was evaluated in live/single Lineage^-^ CD123^+^BDCA4^+^HLADR^+^ gated pDC and depicted for each studied patients or HD together with mean ± SEM values. Surface expression of PD-L1, CD80, CD86 (**D**) and CXCR4, CXCR3, CD62-L, CCR7 (**F**), was determined as mean fluorescence intensity (MFI) and shown results were mean relative values ± SEM of analyzed patients or HD. P-values were depicted as follows: *p≤0.05; ** p≤0.001; *** p≤0.0001.

Further analysis of pDC phenotype in these groups highlighted that CP-AS mainly display a PD-L1^+^ P1 phenotype; while, on the contrary, in CP mainly PD-L1^+^CD80^+^ P2-pDC were observed (**Fig 7B and 7C**). These data were in accordance also to the surface level of both PD-L1 and CD80 markers in pDC expressed in terms of mean fluorescence intensity (MFI) (**Fig 7D** and **S3B Fig**). Importantly, CD86, which was significantly expressed in CP pDC, was further enhanced in CP-AS indicating that asymptomatic infection with SARS-CoV-2 strongly activates pDC (**Fig 7D** and **S3B Fig**).

We then monitored the expression of CXCR3 and CXCR4 on pDC, responsible for their localization to infected peripheral tissues including skin and epithelia, as well as of CCR7 and CD62-L, a chemokine receptor and an integrin that promote pDC lymph node homing respectively (**Fig 7E and 7F**). SARS-CoV-2 natural infection significantly increased both frequency (**Fig 7E**) and surface expression (**Fig 7F** and **S3C Fig**) of all these chemotactic markers on pDC of COVID-19 patients independently of the degree of disease severity, as compared to the level found in circulating pDC of matched HD, indicating that these cells are committed to migrate to the sites of viral infection.

### Asymptomatic and hospitalized individuals with SARS-CoV-2 infection display a specular anti-viral and pro-inflammatory profile

To investigate if the significant difference in number and activation status observed in pDC of asymptomatic and severe COVID-19 patients would mirror a different anti-viral state in these individuals, we analyzed in *ex vivo* PBMC derived from HD, CP-AS and CP the transcription of the classical ISG, Mx1, and found a striking difference in its expression level (**Fig 8A**). RNA from CP-AS PBMC displayed a very high level of this gene as compared to HD, while CP PBMC had a much lower expression, even if significantly higher than HD cells (**Fig 8A**). Interestingly and in accordance with these results, IFN-α level markedly raised in sera of CP-AS as compared to CP and HD (**Fig 8B**).

**8 Fig.**
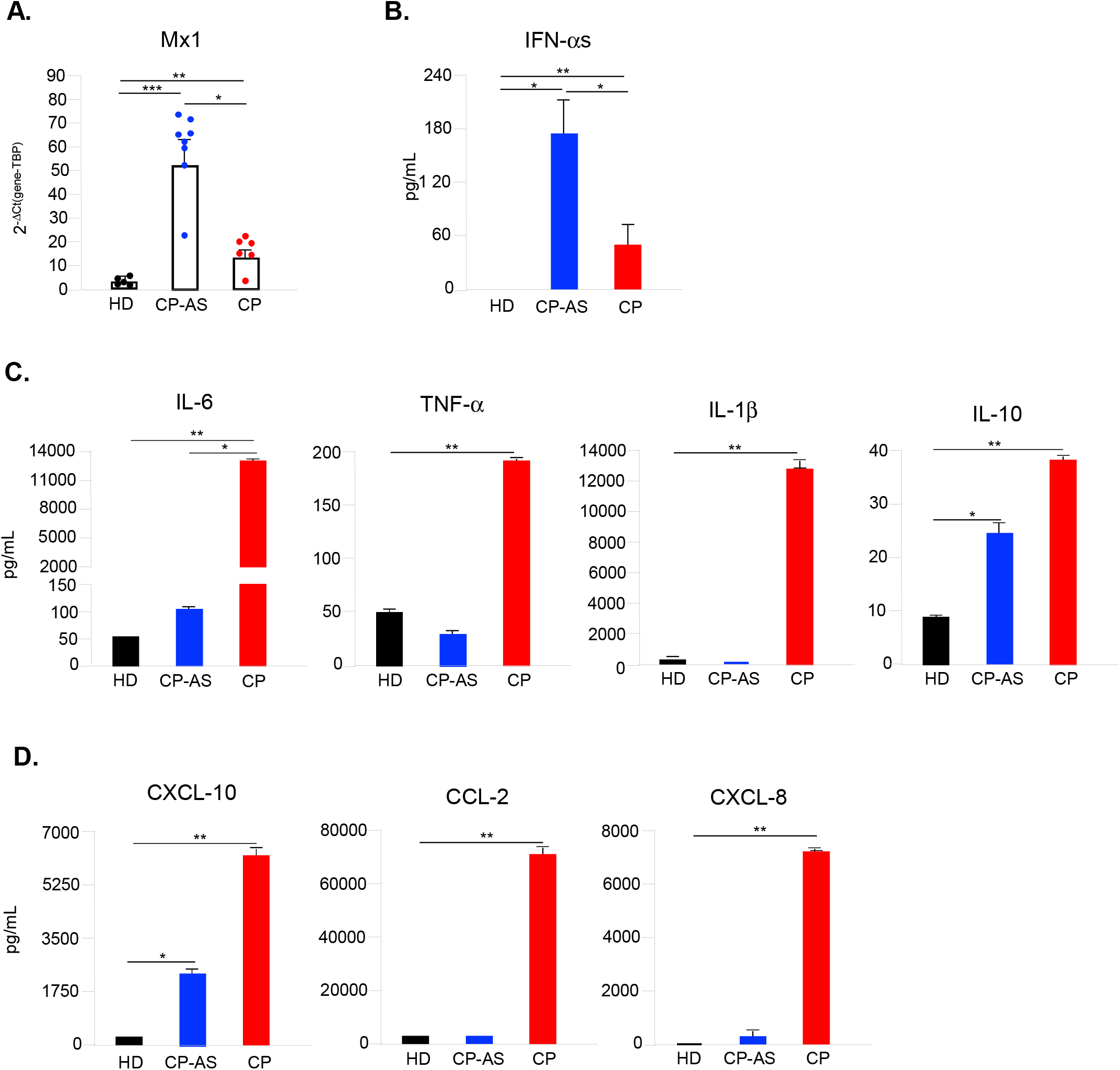
COVID-19 asymptomatic and hospitalized patients display a specular anti-viral and pro-inflammatory profile. Peripheral blood mononuclear cells (PBMC) and sera were collected from asymptomatic (CP-AS, n=8) and hospitalized COVID-19 patients (CP, n=6) as well as matched healthy donors (HD, n=5). (**A**) Relative expression of Mx1 gene was measured by quantitative real time PCR analysis and normalized to TBP level by using the equation 2^−ΔCt^ in total RNA isolated from *ex vivo* PBMC. (**B**) Production of IFN-αs was tested in serum samples by a specific ELISA kit. Production of cytokines (IL-6, TNF-α, IL-1β, IL-10) (**C**) and chemokines (CXCL-10, CCL-2, CXCL-8) (**D**) was tested by multiparametric cytokine bead arrays in collected serum samples. Shown results were mean relative values ± SEM of analyzed patients or HD. P-values were depicted as follows: *p≤0.05; ** p≤0.001; *** p≤0.0001.

The pro-inflammatory cytokines IL-6, TNF-α and IL-1β (**Fig 8C**) together with the chemotactic factors CXCL-10, CCL-2 and CXCL-8 (**Fig 8D**) had a specular profile to that observed for IFN-αs. Indeed, they were all found more increased in sera of critical than in asymptomatic COVID-19 subjects (**Fig 8C and 8D**). Interestingly, the serum level of IL-10, a regulatory cytokine with a strong anti-inflammatory potential also known to be induced by type I IFN, was significantly increased also in CP-AS (**Fig 8C**). Similarly, serum CXCL-10, whose transcription is mainly regulated by type II IFN-γ-mediated signaling but in part also by type I IFNs, was enhanced not only in CP but partly also in CP-AS (**Fig 8D**).

## DISCUSSION

Early phases of anti-viral responses mainly consist of two interconnected pathways: first, the engagement of cellular anti-viral defenses, which are mediated by the transcriptional induction of type I and III IFNs with subsequent up-regulation of ISGs (23) and, second, the recruitment and coordination of specific subset of leukocytes primarily arranged by chemokine secretion (24). In a variety of infection models, including *in vitro* infection of permissive lung epithelium cell lines and primary bronchial epithelial cells as well as *ex vivo* samples derived from COVID-19 patients and animals, it was shown that, despite viral replication, a waning host immune response was induced with an altered induction of type I and III IFNs associated with a high release of chemokines (12, 25). Accordingly, *in vivo* data from severely ill SARS-CoV-2-infected patients confirmed an impaired type I IFN response accompanied by elevated pro-inflammatory cytokine levels in sera (7, 12), thus identifying exacerbated and abnormal responses in the innate branch of the immune system as a main driver of major illness (25).

In the present study, we interrogated human PBMC, a mixture of highly specialized immune cells, to understand their contribution to COVID-19 immunopathogenesis and the dynamic interaction between immune system and SARS-CoV-2 virus. We showed that human PBMC, even if not allowing a productive viral replication, respond to stimulation with a SARS-CoV-2 clinical isolate by expressing high levels of type I and III IFNs and a concomitant ISG transcription, as well as of the inflammatory cytokines IL-6, TNF-α and IL-1β.

It is well known that lymphopenia, together with neutrophilia and monocytopenia, is a characteristic of patients with severe COVID-19 (26). Consistently with these data, one of the main mechanisms associated with disease severity is the recruitment of inflammatory immune cells towards infected lungs and the subsequent hyperinflammatory state or the so-called cytokine releasing syndrome, which includes inflammatory mediators as well as chemokines (6, 27). A dramatic induction of chemotactic factors was found in *post-mortem* lung samples and in plasma or sera of COVID-19 patients at different stages of disease (28). In particular, levels of CXCL-10, involved in the etiology of various pulmonary conditions and attracting monocytes, NK and CXCR3-expressing Th1 cells (29, 30), increased with disease severity, suggesting that it may help in early diagnosis and could serve as a potential predictive marker of disease outcome (27). Also, the monocyte chemoattractant molecule CCL-2, was found upregulated during the early phase of infection, further increased during late stages of fatal disease (31) and associated with a prolonged duration of intensive care unit stay (32, 33). Patients with fatal COVID-19 also showed significantly higher plasma levels of the neutrophil recruiting CXCL-8 (34) as well as of the T and NK cell-attracting CXCL-9 (35), as compared to severe and/or mild COVID-19 patients (12, 31). In our study, *in vitro* SARS-CoV-2 stimulation of PBMC recapitulates the *in vivo* scenario found in COVID-19 since we observed a strong release of the aforementioned chemokines emerged as crucial in COVID-19 pathogenesis.

Further, we proved that SARS-CoV-2-mediated release of inflammatory cytokines and chemokines heavily relies on the TLR-7/8 signaling and requires stimulation with infectious live SARS-CoV-2 virions, consistently with recently published evidences of SARS-CoV2 partially productive infection of lymphomononuclear cell subsets (21, 36).

A characteristic of severe COVID-19 cases and of *in vitro* infected SARS-CoV-2 permissive respiratory epithelial cells is the highly impaired type I IFN release and response (with no IFN-β and low IFN-α production and activity detected), thus allowing and sustaining viral replication and an exacerbated inflammatory response (7, 12). With our experiment on permissive Calu-3 lung epithelial cells conditioned with culture supernatants of SARS-CoV-2-stimulated PBMC, we proved, for the first time, that PBMC-derived type I and III IFNs are bioactive and strongly enhanced ISG transcription with a concomitant anti-viral state in Calu-3 cells, differently to what found in the SARS-CoV-2 infected counterpart. Consistently with these data, SARS-CoV-2-stimulated PBMC supernatant partially inhibits SARS-CoV-2 replication in Calu-3 infected cells. Importantly, our results are fully coherent with previous data showing that *in vitro* administration of type I IFN-β (37) and type III IFN-λ (38) restrains and reduces SARS-CoV-2 viral replication.

Type I IFNs emerged as key protective factors in preventing COVID-19 disease severity. Genetical analyses of COVID-19 patients with life-threatening pneumonia had, indeed, reported inborn errors of type I IFN immunity and variants in genes encoding for RNA sensors and type I IFN regulating elements in absence of other risk factors (39, 40). Furthermore, more than 10% of COVID-19 patients with severe pneumonia had neutralizing auto-antibodies against either type I IFN-ϖ or IFN-αs, or both, before disease onset and 94% of them were males; findings that might also explain the excess of men among the individuals with life-threatening COVID-19 disease (41). Moreover, severe COVID-19 patients uniquely produce antibodies that functionally block, through binding of the Fc domain to FcγR-IIb, the production of type I IFN and ISG-expressing cells found in mild disease (25).

Having in mind all these evidences, we investigated which immune cell type is responsible for type I IFN release upon SARS-CoV-2 stimulation. pDC are known to be the principal cell type among immune cells specialized in type I IFN release sensing viral RNA in a TLR7-dependent manner (19). pDC can sense almost all viruses, including the coronaviruses SARS-CoV and Middle East respiratory syndrome coronavirus (MERS) (42–44), and were shown to recognize MERS *via* TLR7 (44). In our study, pDC isolated from peripheral blood of healthy donors and stimulated *in vitro* with SARS-CoV-2 produced a high amount of TLR7-mediated IFN-αs, with a level comparable to that found in PBMC cultures, even if pDC do not express both the entry factors ACE-2 or TMPRSS2 [our data and (45)].

Further, our study showed that stimulation with SARS-CoV-2 regulated pDC phenotype and activation status. Hence, viral sensing mediates the cell surface up-regulation of PD-L1 on more of the 55% of cultured pDC, a phenotype found in pDC specialized in type I IFN production (22). In presence of a TLR-7/8 inhibitor SARS-CoV-2-treated pDC turn from PD-L1^+^CD80^−^ to PD-L1^+^CD80^+^ or CD80^+^ only expressing cells, two subpopulations known to mediate adaptive functions and T cell interaction (22), as exemplified in our study also by the up-regulation of the costimulatory and maturation-associated marker CD86. Accordingly, pDC in presence of SARS-CoV-2 and TLR7/8 inhibitor do not release type I IFNs and turn into TNF-α- and IL-6-producing cells. Production of the pro-inflammatory cytokine TNF-α, in particular, is known to negatively control in a autocrine/paracrine fashion the production of IFN-αs in pDC as soon as the maturation process starts (46). While we were finalizing this paper, Onodi and colleagues published similar results showing that SARS-CoV-2, in absence of productive infection, induces pDC phenotype diversification at a level similar to influenza virus (45). SARS-CoV-2-mediated type I IFN production was then blocked in pDC by hydroxychloroquine (45), in line to what we found in presence of a specific TLR7/8 inhibitor.

SARS-CoV-2 rapidly impairs T cell and DC responses during the acute phase of infection, which could have significant implications for COVID-19 pathogenesis (47). In particular, pDC have been extensively studied in patients with severe COVID-19 and were found significantly reduced or absent in the blood of these individuals (47–49). Importantly, while inflammatory monocytes and CD1c^+^ conventional DC were present in lung infiltrates of patients with severe COVID-19, CD123^hi^ pDC were depleted in blood but absent in the lungs (50).

While Onodi *et al.* demonstrated *in vitro* that pDC activation by SARS-CoV-2 is dependent on IRAK4- and UNC93B1-mediated signaling pathways by means of patients genetically deficient in genes encoding these factors (45), in this study we deepen the current understanding on pDC contribution and phenotype diversification in SARS-CoV-2 infection by characterizing *ex vivo* blood samples derived from patients at different COVID-19 disease severity. As expected, we observed a profound pDC depletion, in terms of both frequency and absolute numbers, in patients hospitalized with severe COVID-19 as compared to matched HD. Interestingly, then, in a cohort of asymptomatic SARS-CoV-2-infected individuals circulating pDC were already reduced in the peripheral blood as respect to the healthy counterpart but significantly higher than what found in hospitalized COVID-19 patients. Nonetheless, the phenotype was drastically different in pDC derived from severely infected or asymptomatic subjects. pDC from asymptomatic COVID-19 mainly expressed PD-L1, while cells derived from severe COVID-19 were consistently represented by PD-L1^+^CD80^+^ phenotype. Importantly, the analysis of costimulatory marker CD86, that was significantly expressed in pDC from critically ill patients and further enhanced in cells of asymptomatic subjects, indicated that pDC are strongly activated during asymptomatic infection.

We also observed that the expression of chemokine receptors responsible for pDC localization to infected peripheral tissues, such as CXCR3 and CXCR4, or for their lymph node homing, as CCR7 and CD62-L, was induced by SARS-CoV-2 natural infection both in terms of frequency and surface expression levels in cells of both asymptomatic or severe COVID-19 patients, possibly indicating that these cells are committed to migrate to the sites of viral infection independently of the degree of disease severity.

Hence, having found dramatic differences in phenotype of pDC from asymptomatic or severe COVID-19, we also tested if these data matched to the induced cytokine profile in the different disease courses. Most interestingly, PBMC isolated *ex vivo* from HD, asymptomatic or severely infected subjects indicated that anti-viral ISG expression is massively induced only in asymptomatic individuals. These data matched to the high amount of IFN-αs present in sera and with the PD-L1^+^ P1 phenotype, specialized in type I IFN production, found in pDC of asymptomatic SARS-CoV-2 infected subjects. These results are in line with recent findings proving the high frequency of neutrophils and monocytes expressing high levels of ISG only in mild COVID-19, and not in the severe form (25). These ISG are likely up-regulated in these cell types in response to the high amount of circulating IFN-αs released by PD-L1^+^ pDC.

Lastly, our study also demonstrated that a specular profile of anti-viral and inflammatory cytokines and chemokines exists between sera samples derived from asymptomatic or severe SARS-CoV-2 infection. We found that severe COVID-19 patients displayed, as extensively already reported (35, 51, 52), high level of circulating inflammatory cytokines, including IL-6, TNF-α, IL-1β and chemokines such as CXCL-10, CCL-2 and CXCL-8 contributing to the diffuse inflammatory status and the so-called cytokine storm. Conversely, we demonstrated that subjects with asymptomatic infection have high level of IFN-αs and of the immune-regulatory IFN-inducible IL-10 known to dampen ongoing inflammatory responses.

In conclusion, in our study by using an *in vitro* human PBMC-based experimental model we recapitulated the *in vivo* scenario found in early SARS-CoV-2 infection and assessed the importance of pDC response and pDC-induced type I IFN in the regulation of the anti-viral state in asymptomatic and severe COVID-19 patients. Thus, the PBMC-based experimental setting might represent an optimal tool to study SARS-CoV-2-induced immune responses. Modulating innate antiviral immunity and, in particular, the pDC/type I IFN axis since the early phases of COVID-19 may help to pinpoint novel pharmacological strategies or host-directed therapies that would counter-act the raising of hyper-inflammation and the resulting diffuse damage contributing to a rapid resolution of SARS-CoV-2 infection.

## MATERIALS AND METHODS

### Patients

Istituto Superiore di Sanità Review Board approved the use of blood from HD (AOO-ISS - 14/06/2020 - 0020932) and from asymptomatic SARS-CoV2 infected individuals (IN_COVID, AOO-ISS - 22/03/2021 - 0010979). Policlinico Tor Vergata approved blood withdrawal from hospitalized COVID-19 patients (COVID-SEET, CE#46.20, 18/04/2020).

In particular, for this study six hospitalized COVID-19 [CP; 2 females/4 males; median age ±Standard Deviation (SD) 51.5 ± 23.3 yrs.] and eight asymptomatic [CP-AS; 4 females/4 males; 58.5 ± 10.4 yrs.] patients matched to five HD [2 females/3 males; 53 ± 12.5 yrs.] were enrolled and provided written informed consent. Main demographic, clinical and experimental data related to asymptomatic and hospitalized COVID-19 patients are listed in **S2** and **S3 Tables**.

### Isolation and culture of PBMC, pDC and monocytes

PBMC were collected from peripheral blood and isolated and cultured as described (53). pDC and monocytes were purified from isolated PBMC by magnetic separation by using anti-BDCA4 and anti-CD14 microbeads (Miltenyi biotech) respectively, as previously described (54). The purity of the recovered cells was greater than 95% as assessed by flow cytometry analysis with anti-BDCA4 (Miltenyi biotech) or anti-CD14 (BD Biosciences) monoclonal antibodies.

### Virus production

Vero E6 (Vero C1008, clone E6-CRL-1586; ATCC) cells were cultured in Dulbecco’s Modified Eagle Medium (DMEM) supplemented with non-essential amino acids (NEAA, 1x), penicillin/streptomycin (P/S, 100 U/mL), HEPES buffer (10 mM) and 10% (v/v) Fetal bovine serum (FBS). A clinical isolate hCoV-19/Italy/UniSR1/2020 (GISAID accession ID: EPI_ISL_413489) was isolated and propagated in Vero E6 cells, and viral titer was determined by 50% tissue culture infective dose (TCID_50_) and plaque assay for confirming the obtained titer.

### Virus Titration

Virus stocks and supernatants of the experimental conditions were titrated using Endpoint Dilutions Assay (EDA, TCID_50_/mL). Vero E6 cells (4 × 10^5^ cells/mL) were seeded into 96 wells plates and infected with base 10 dilutions of collected medium, each condition tested in triplicate. After 1 h of adsorption at 37°C, the cell-free virus was removed, and complete medium was added to cells. After 72 h, cells were observed to evaluate CPE. TCID_50_/mL was calculated according to the Reed–Muench method.

### Viral inactivation by UV-C

An aliquot (0.170 mL) of viral stock was place in a 24-well plate in ice to counteract irradiation-derived heating of the sample and irradiated with approximately 1.8 mW/cm^2^ at a work distance of 20 cm for 40 minutes. The viral inactivation was checked using undiluted supernatant in an infection assay.

### Calu-3 treatment and infection

Calu-3 (Human lung cancer cell line, ATCC HTB-55) were cultured in MEM supplemented with NEAA (1x), P/S (100 U/mL), Sodium Pyruvate (1 mM), and 10% (v/v) FBS. Calu-3 cells were seeded into 12-wells plates to reach confluence. Before the viral adsorption, the cells were treated with 200 μL of the different PBMC supernatants for 1 hour at 37°C. The viral adsorption was conducted as already described using 5.62 × 10^4^ TCID_50_/mL of hCoV-19/Italy/UniSR1/2020. After then, 500 μL of the different PBMC supernatants were added and the cells were incubated at 37°C for 24 hours.

### Cell stimulation and supernatant collection

PBMC, pDC and monocytes were pre-incubated for 1 hour at 37°C with infectious or UV-inactivated SARS-CoV2 at 0.01, 0.02, 0.04 or 0.1 MOI and then cultured at 2×10^6^ cells/ml in RPMI 1640 in presence of P/S (100 U/mL), L-glutamine (2mM) and 10% FBS for 24 or 48 hours. As mock-treatment, cells were stimulated with supernatants from Vero E6 uninfected cells at a dilution corresponding to that of MOI 0.1 infected cultures. Cells were also treated with R848, a specific TLR7/8 agonist (5μM, Invivogen) as positive control, or with 1000 U/mL of recombinant IFN-α2 and IFN-β (Peprotech). Furthermore, where indicated, PBMC and pDC were also pre-treated for 30 min with 1μM TLR7/8 antagonist, 2087c oligonucleotide (Miltenyi biotech), prior to stimulation with SARS-CoV2.

After 24 or 48 hours, cell culture supernatants were harvested and treated for 30 minutes at 56°C, then stored at −80°C for later use. SARS-CoV2 inactivation was tested by back titration for each experiment (see above for details).

### Detection of cytokines and chemokines in culture supernatants

Release of IFN-αs was measured by a specific ELISA kit (PBL assay science). Production of cytokines (IL-6, TNF-α, IL-1β, IL-10 and IL-12p70) and chemokines (CXCL-10, CCL-2, CXCL-9, CXCL-8 and CCL-5) was quantified by specific cytometric bead arrays (BD Biosciences) on a FACS Canto (BD Biosciences) and analyzed by FCAP array software (BD Biosciences).

### Flow cytometry analyses

Monoclonal antibodies anti-Lineage cocktail (Lin), PD-L1, CD80, CD86, HLA-DR, CD123, CXCR-3, CXCR-4, CD62-L and CCR7 as well as IgG1 or IgG2a isotype controls were purchased from BD Biosciences, while BDCA4 from Miltenyi Biotech. To establish cell viability and exclude dead cells from flow cytometry analyses, Fixable Viability Dye (FvDye, eBioscience) was always included in antibody cocktails. In the mixed cell population of PBMC, pDC were considered as those cells in live/single FvDye^-^Lineage-CD123^+^BDCA4^+^HLADR^+^ gate (**S3A Fig**). Cells (1.5×10^6^ for PBMC or 5×10^4^ for isolated pDC) were incubated with monoclonal antibodies at 4°C for 30 min and then fixed with 4% paraformaldehyde before analysis on a Cytoflex cytometer (Beckman Coulter). Data were analyzed by Flow Jo software v.10.7 (BD Biosciences). Expression of analyzed cell surface molecules was evaluated using the median fluorescence intensity (MFI). Only viable and single cells were considered for further analysis.

### RNA isolation and quantitative real time PCR analysis

Total RNA was isolated by Trizol® Reagent (Invitrogen, Thermo Fischer Scientific), quantified using a Nanodrop2000 spectrophotometer and quality assessed with an established cut-off of ~1.8 for 260/280 absorbance ratio. Reverse-transcription was conducted by Vilo™ reverse transcriptase kit (Invitrogen, Thermo Fischer Scientific).

Expression of genes encoding Mx1, IFN-αs, IFN-β, IFN-λ1, IL-6, ACE-2 and TMPRSS2 was measured by quantitative real time PCR (q-PCR) using the appropriate TaqMan™ assay and TaqMan™ Universal Master Mix II (Applied Biosystems, Thermo Fisher Scientific) on a ViiA™ 7 Instrument (Applied Biosystems, Thermo Fisher Scientific).

The housekeeping gene TATA-box-binding protein (TBP) was used as normalizer. Real time reactions were run at least in duplicates. Sample values for each mRNA were normalized to the selected housekeeping gene using the formula 2^−ΔCt^.

## Supporting information

Severa_Suppl_Figures

## Statistical analysis

Statistical analysis was performed using One-way Repeated-Measures ANOVA when three or more stimulation conditions were compared. A two-tailed paired Student’s t-test was used when only two stimulation conditions were compared. Results were shown as mean values ± SEM. P ≤0.05 was considered statistically significant. In the figures, star scale was assigned as follows: *=p ≤ 0.05; **=p ≤ 0.01; ***=p ≤ 0.001.

## ACKNOWLEDGEMENTS

The authors acknowledge Dr. Valentina Tirelli and Dr. Massimo Sanchez of Flow Cytometry Facility (FAST, Istituto Superiore di Sanità, Rome, Italy), Concetta Fabiani, Eleonora Benedetti and Angela Di Martino (Department of Infectious Diseases, Istituto Superiore di Sanità, Rome, Italy), and Dr. Silvia Riccetti (Department of Molecular Medicine, University of Padua, Padua, Italy) for technical support.

This work was funded by Istituto Superiore di Sanità and partly co-financed by the Italian Ministry of Health (grant GR-2016-02363749 to MS) and the European Union’s Horizon 2020 research and innovation programme, under grant agreement no. 874735 (VEO) to LB.

## DATA AND MATERIALS AVAILABILITY

This study did not generate new unique reagents/dataset/code. All data are available in the main text or the supplementary materials.

## SUPPORTING INFORMATION

**S1 Fig. Kinetic of expression of cytokine and chemokine in PBMC stimulated by SARS-CoV-2.**Peripheral blood mononuclear cells (PBMC) were left untreated (not stimulated, ns) or stimulated for 24 or 48 hours with SARS-CoV-2 (MOI=0.04 and MOI=0.1) (**A, B**) Production of cytokine IL-6 (**A**) and chemokines CXCL-10 and CXCL-9 (**B**) was tested by multiparametric cytokine bead arrays in collected cell culture supernatants. Shown results were mean relative values ± SEM of 3 independent experiments. P-values were calculated by two-tailed Students’ t-test and were depicted as follows: *p≤0.05; **p≤0.01.**S2 Fig.** Analysis of SARS-CoV-2 entry factors in pDC and monocytes and cytokine production in isolated CD14+ monocytes.

**S2 Fig. Analysis of SARS-CoV-2 entry factors in pDC and monocytes and cytokine production in isolated CD14+ monocytes.**(**A, B**) pDC, monocytes and Calu-3 cell line were left not stimulated (ns) or stimulated with 1000 U/mL of recombinant IFN-α or IFN-β. Gene expression of ACE-2, TMPRSS2 and Mx1 was measured by quantitative RT-PCR. (**C**) Monocytes were left untreated (not stimulated, ns) or stimulated for 24 hours with SARS-CoV-2 (MOI=0.04 and MOI=0.1). Production of IFN-αs and IL-6 was tested by specific ELISA in collected cell culture supernatants. Shown results were mean relative values ± SEM of 3 independent experiments.

**S3 Fig. Representative gating strategy for pDC phenotypical analysis in COVID-19 patients.** Freshly isolated peripheral blood mononuclear cells (PBMC) from asymptomatic (CP-AS, n=8) and hospitalized COVID-19 patients (CP, n=6) and matched healthy donors (HD, n=5) were stained with a well-established antibody cocktails to study by flow cytometry plasmacytoid dendritic cell (pDC) diversification and activation status as well as expression of chemokine receptors. (**A**) The percentage (%) of total pDC was evaluated in Fixable viability dye (FvDye)^−^ live Lineage-CD123^+^BDCA4^+^ gated cells. In pDC we also evaluated the % of PD-L1 or CD80 expressing cells. (**B, C**) Surface expression of PD-L1, CD80, CD86 (**B**) and CXCR4, CXCR3, CCR7, CD62-L (**C**) was determined as mean fluorescence intensity (MFI) in CP or HD gated pDC. Shown dot plots and histograms are representative of all analyzed CP and HD.

**S1 Table. Evaluation of cell death in human PBMC cultures treated with live SARS-CoV2.**

**S2 Table. Main demographic and clinical characteristics of hospitalized COVID-19 patients.**

**S3 Table. Main demographic and clinical characteristics of asymptomatic COVID-19 patients.**

