## Supplementary material for "Differential plasmacytoid dendritic cell phenotype and type I Interferon response in asymptomatic and severe COVID-19 infection": Severa_Suppl_Figures

### Slide 1
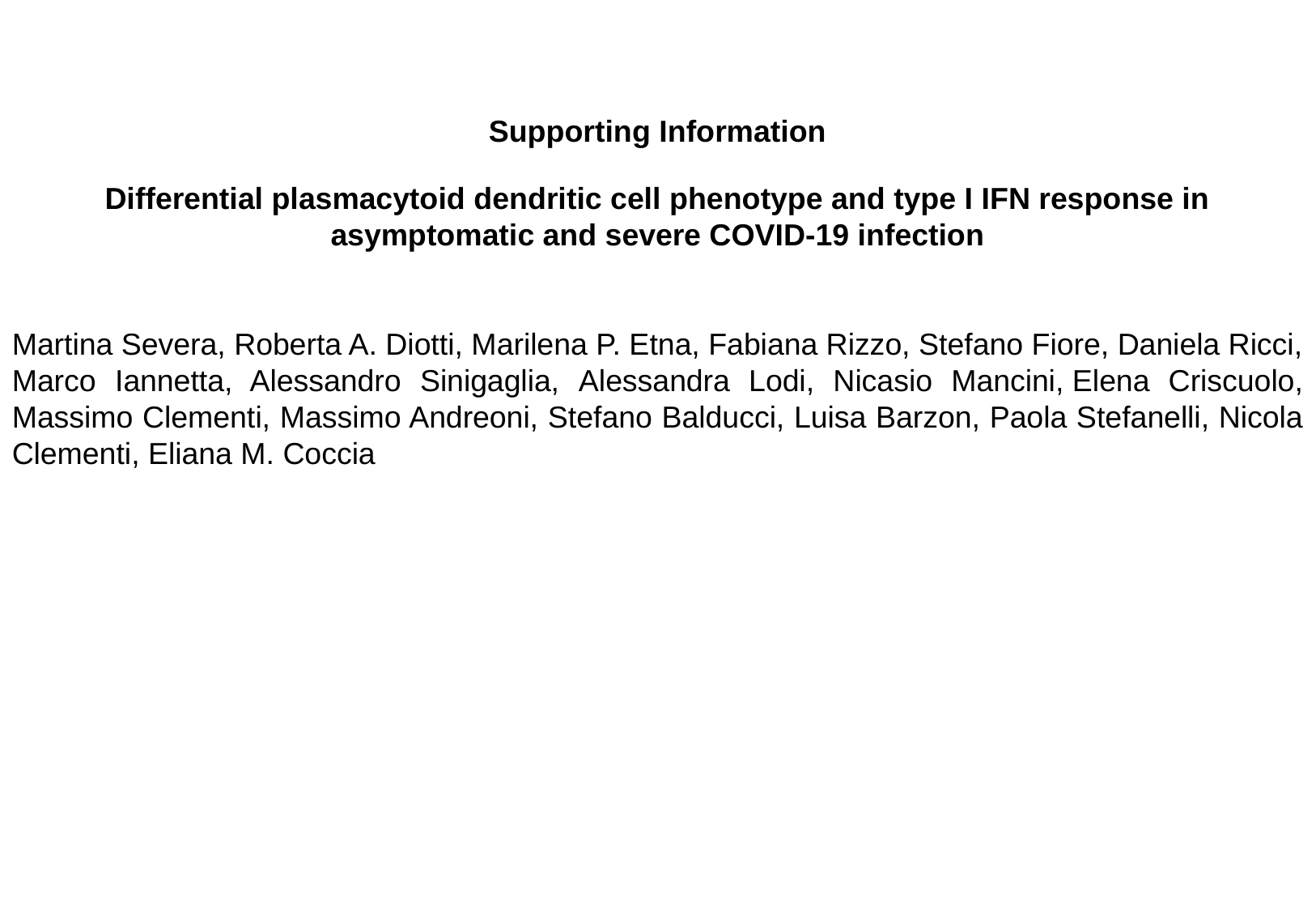

Supporting Information
Differential plasmacytoid dendritic cell phenotype and type I IFN response in asymptomatic and severe COVID-19 infection
Martina Severa, Roberta A. Diotti, Marilena P. Etna, Fabiana Rizzo, Stefano Fiore, Daniela Ricci, Marco Iannetta, Alessandro Sinigaglia, Alessandra Lodi, Nicasio Mancini, Elena Criscuolo, Massimo Clementi, Massimo Andreoni, Stefano Balducci, Luisa Barzon, Paola Stefanelli, Nicola Clementi, Eliana M. Coccia

### Slide 2
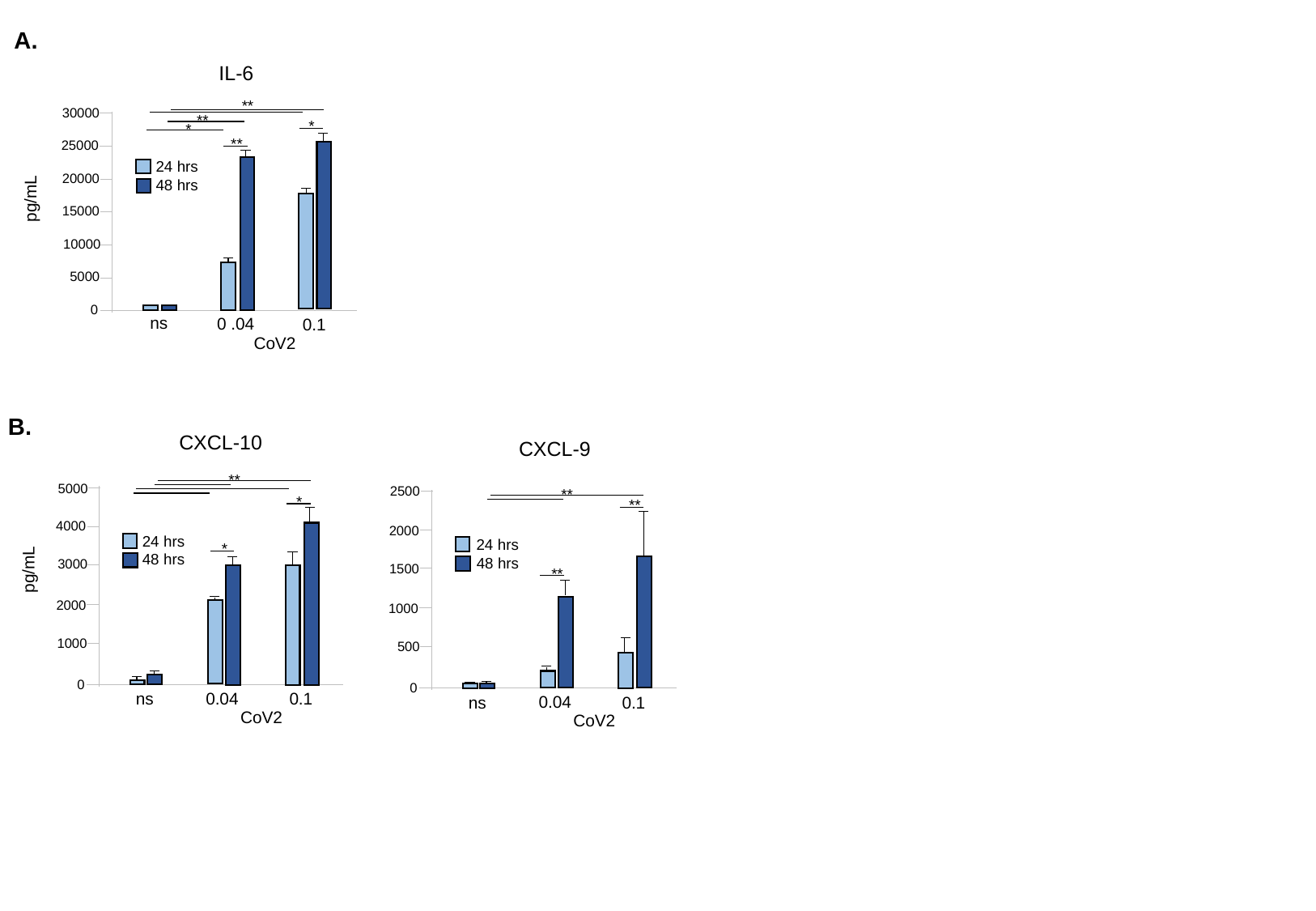

A.
IL-6
**
30000
**
*
*
**
25000
24 hrs
20000
48 hrs
pg/mL
15000
10000
5000
0
ns
0 .04
0.1
CoV2
B.
CXCL-10
**
5000
*
4000
24 hrs
*
48 hrs
3000
pg/mL
2000
1000
0
ns
0.04
0.1
CoV2
CXCL-9
2500
**
**
2000
24 hrs
48 hrs
1500
**
1000
500
0
ns
0.04
0.1
CoV2

### Slide 3
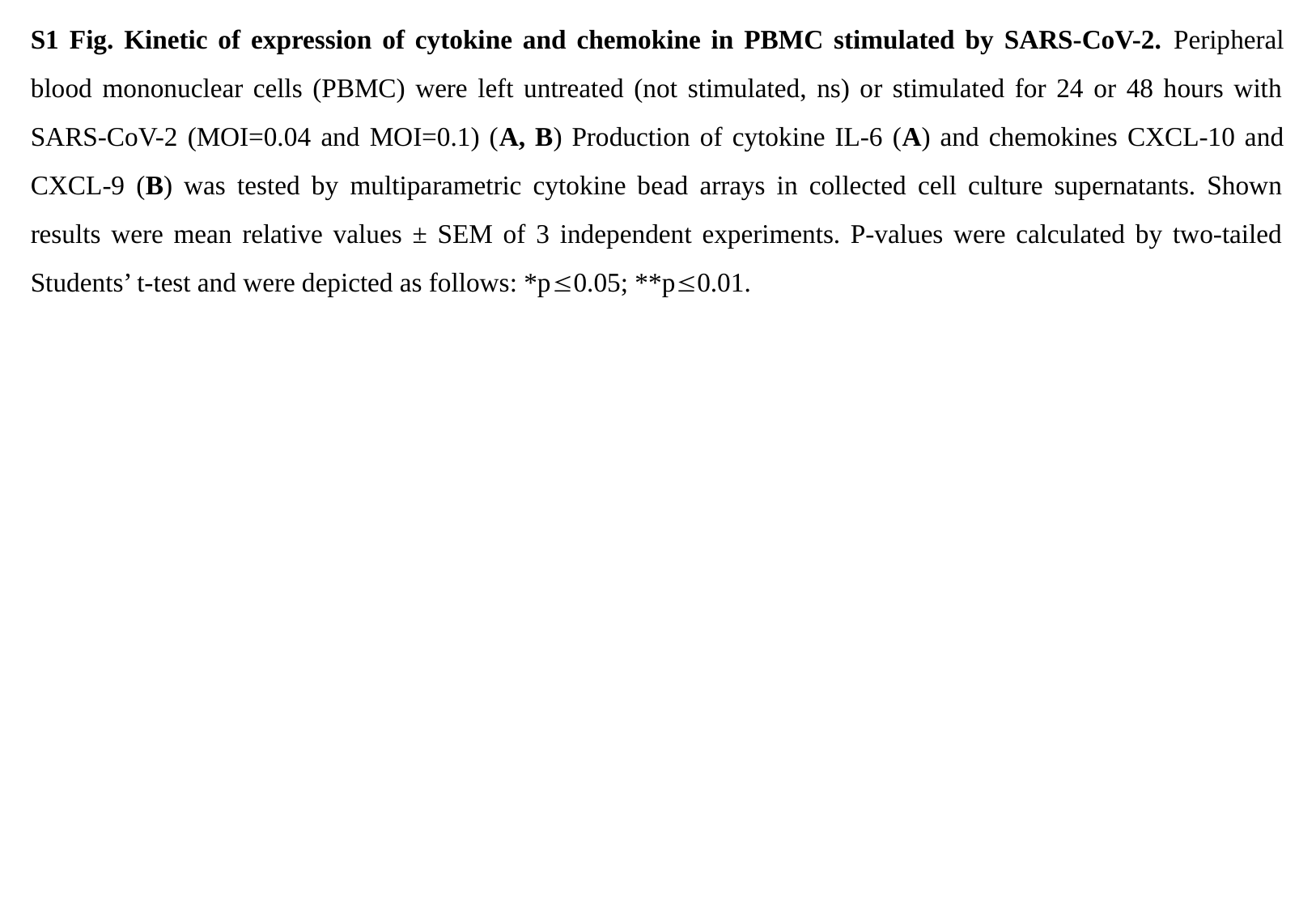

S1 Fig. Kinetic of expression of cytokine and chemokine in PBMC stimulated by SARS-CoV-2. Peripheral blood mononuclear cells (PBMC) were left untreated (not stimulated, ns) or stimulated for 24 or 48 hours with SARS-CoV-2 (MOI=0.04 and MOI=0.1) (A, B) Production of cytokine IL-6 (A) and chemokines CXCL-10 and CXCL-9 (B) was tested by multiparametric cytokine bead arrays in collected cell culture supernatants. Shown results were mean relative values ± SEM of 3 independent experiments. P-values were calculated by two-tailed Students’ t-test and were depicted as follows: *p≤0.05; **p≤0.01.

### Slide 4
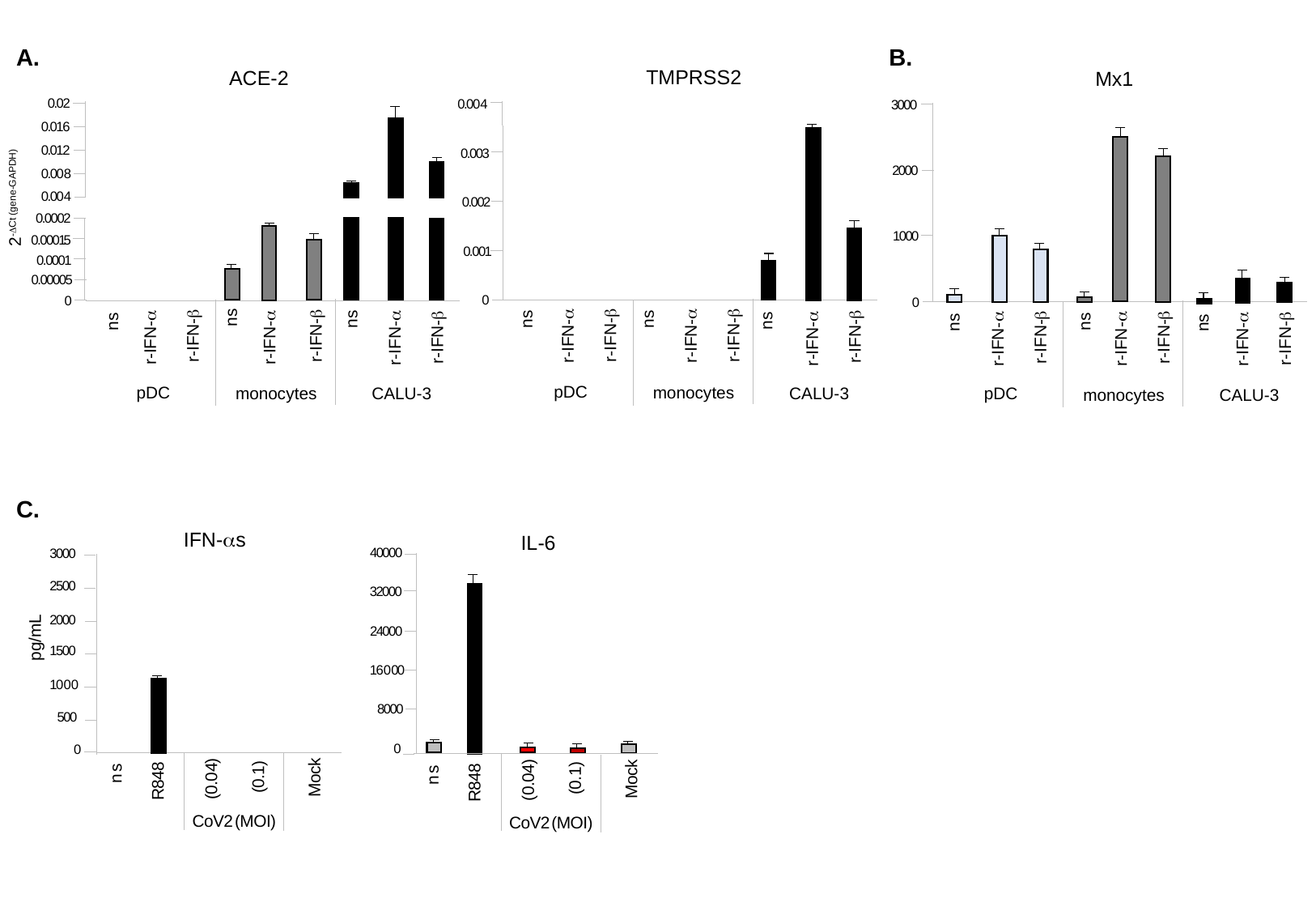

A.
B.
TMPRSS2
0 . 0 0 4
0 . 0 0 3
0 . 0 0 2
0 . 0 0 1
0
ns
ns
ns
r-IFN-β
r-IFN-β
r-IFN-α
r-IFN-α
r-IFN-β
r-IFN-α
pDC
monocytes
CALU-3
ACE-2
0 . 0 2
0 . 0 1 6
0 . 0 1 2
0 . 0 0 8
2^-ΔCt (gene-GAPDH)^
0 . 0 0 4
0 . 0 0 0 2
0 . 0 0 0 15
0 . 0 0 0 1
0 . 0 0 0 0 5
0
ns
ns
ns
r-IFN-β
r-IFN-β
r-IFN-β
r-IFN-α
r-IFN-α
r-IFN-α
pDC
monocytes
CALU-3
Mx1
3 0 0 0
2 0 0 0
1 0 0 0
0
ns
ns
ns
r-IFN-β
r-IFN-β
r-IFN-β
r-IFN-α
r-IFN-α
r-IFN-α
pDC
monocytes
CALU-3
C.
IFN-αs
3 0 0 0
2 5 0 0
2 0 0 0
pg/mL
1 5 0 0
1 0 0 0
5 0 0
0
n s
M o c k
( 0 . 1 )
R 8 4 8
( 0 . 0 4 )
C o V 2 ( M O I )
IL-6
4 0 0 0 0
3 2 0 0 0
2 4 0 0 0
1 6 0 0 0
8 0 0 0
0
n s
M o c k
( 0 . 1 )
R 8 4 8
( 0 . 0 4 )
C o V 2 ( M O I )

### Slide 5
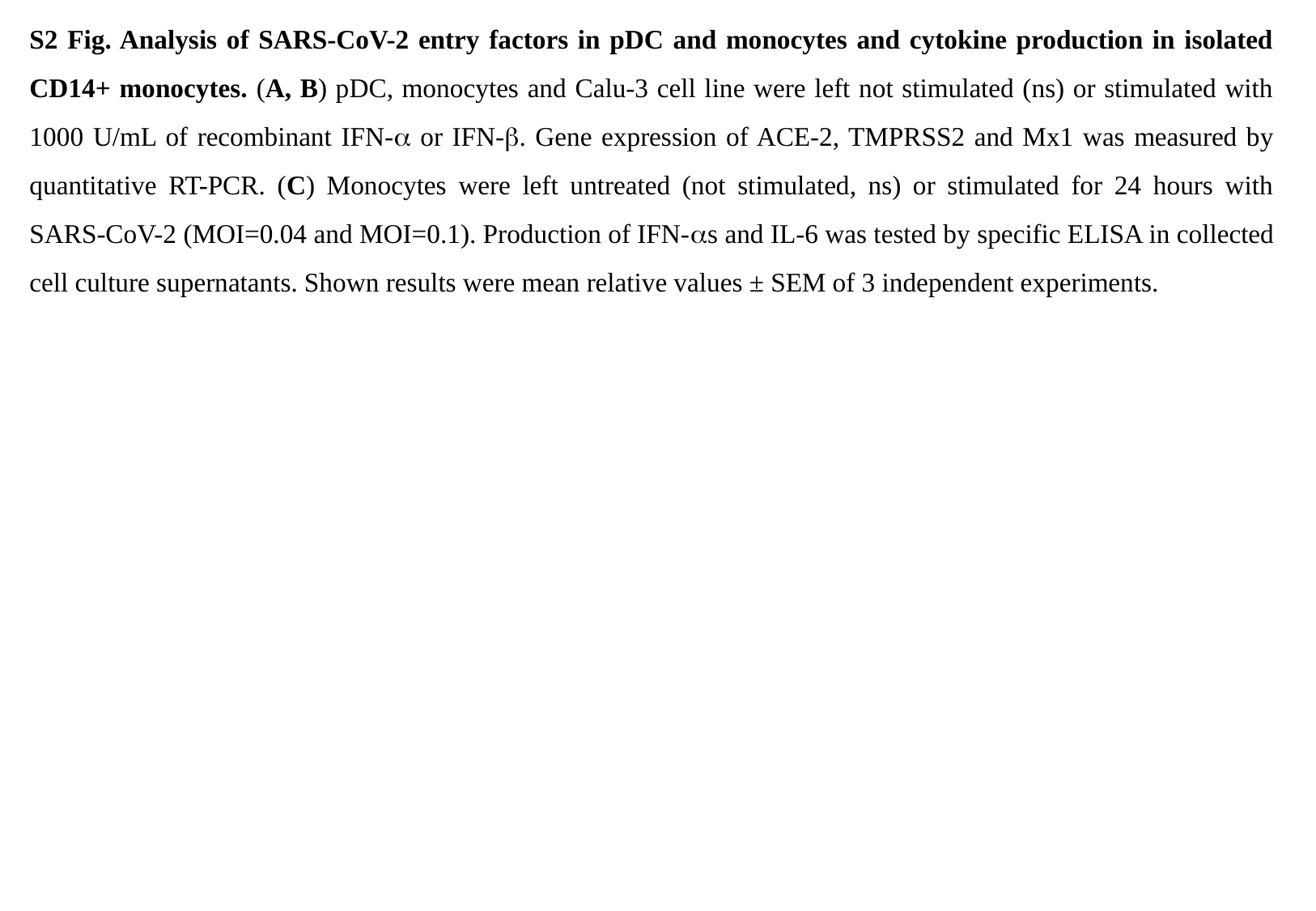

S2 Fig. Analysis of SARS-CoV-2 entry factors in pDC and monocytes and cytokine production in isolated CD14+ monocytes. (A, B) pDC, monocytes and Calu-3 cell line were left not stimulated (ns) or stimulated with 1000 U/mL of recombinant IFN-α or IFN-β. Gene expression of ACE-2, TMPRSS2 and Mx1 was measured by quantitative RT-PCR. (C) Monocytes were left untreated (not stimulated, ns) or stimulated for 24 hours with SARS-CoV-2 (MOI=0.04 and MOI=0.1). Production of IFN-αs and IL-6 was tested by specific ELISA in collected cell culture supernatants. Shown results were mean relative values ± SEM of 3 independent experiments.

### Slide 6
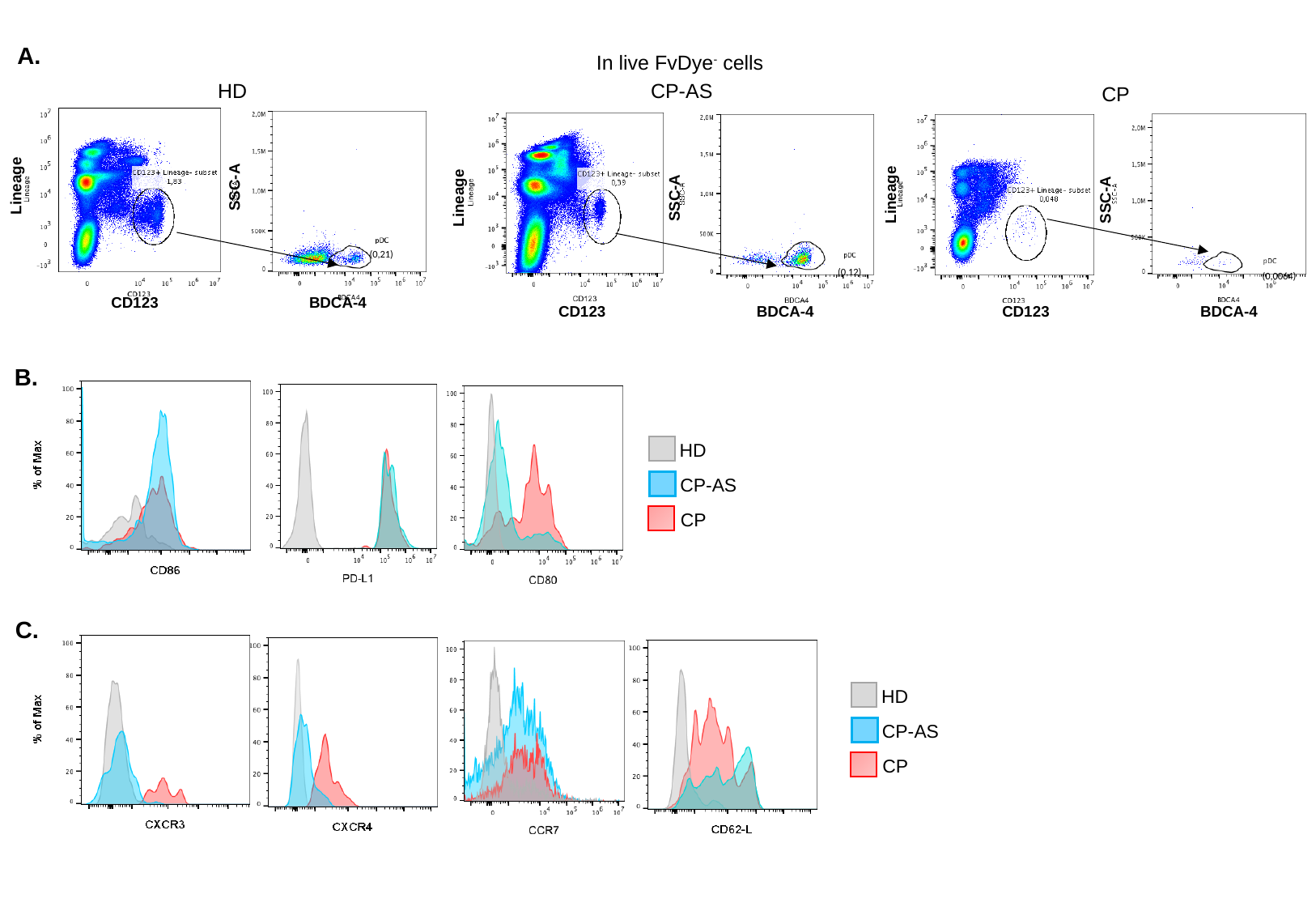

A.
In live FvDye^-^ cells
HD
CP-AS
CP
(0,21)
(0,0064)
(0,12)
Lineage
SSC-A
Lineage
Lineage
SSC-A
SSC-A
CD123
BDCA-4
CD123
BDCA-4
CD123
BDCA-4
B.
HD
CP-AS
CP
C.
HD
CP-AS
CP

### Slide 7
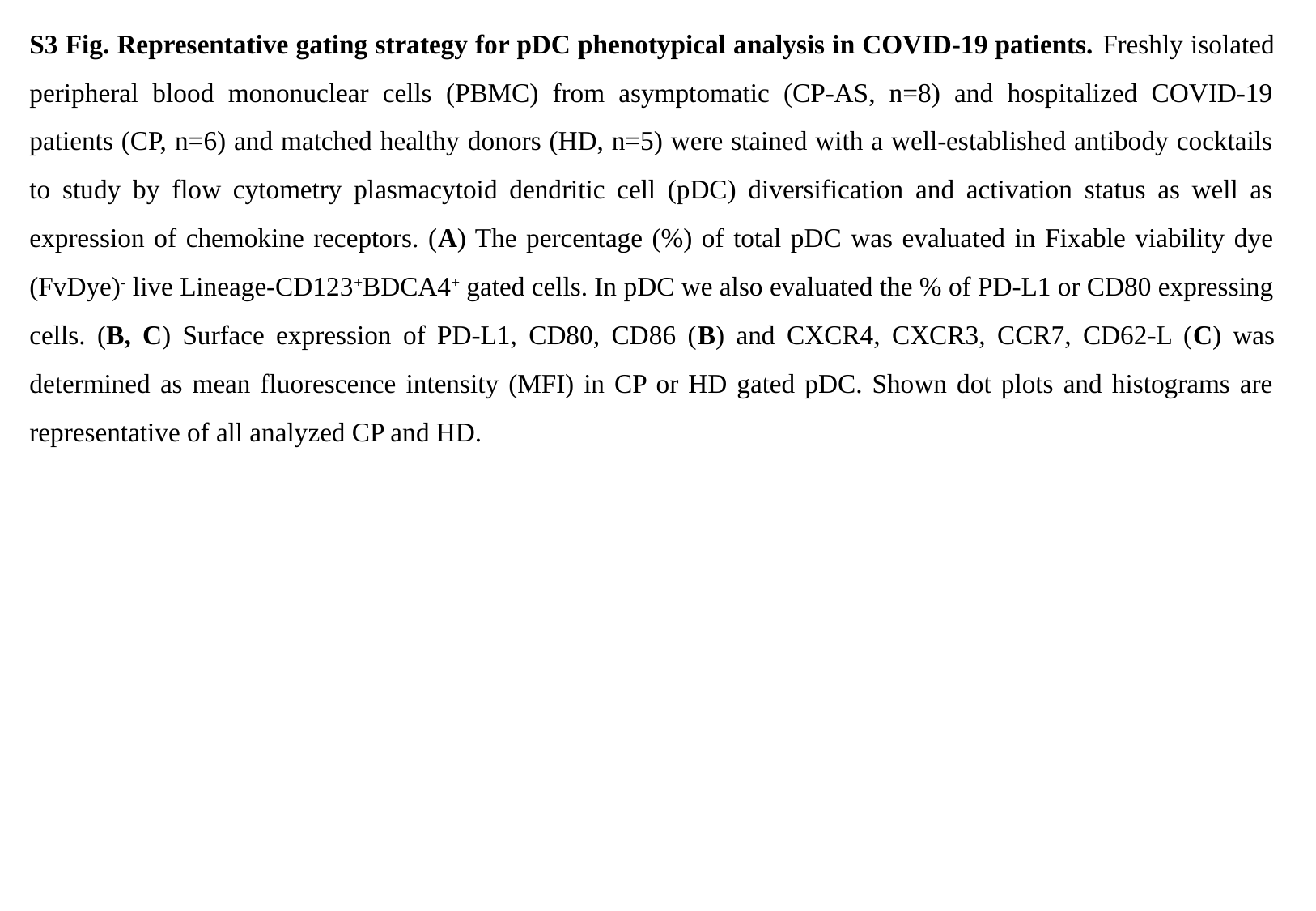

S3 Fig. Representative gating strategy for pDC phenotypical analysis in COVID-19 patients. Freshly isolated peripheral blood mononuclear cells (PBMC) from asymptomatic (CP-AS, n=8) and hospitalized COVID-19 patients (CP, n=6) and matched healthy donors (HD, n=5) were stained with a well-established antibody cocktails to study by flow cytometry plasmacytoid dendritic cell (pDC) diversification and activation status as well as expression of chemokine receptors. (A) The percentage (%) of total pDC was evaluated in Fixable viability dye (FvDye)^-^ live Lineage-CD123^+^BDCA4^+^ gated cells. In pDC we also evaluated the % of PD-L1 or CD80 expressing cells. (B, C) Surface expression of PD-L1, CD80, CD86 (B) and CXCR4, CXCR3, CCR7, CD62-L (C) was determined as mean fluorescence intensity (MFI) in CP or HD gated pDC. Shown dot plots and histograms are representative of all analyzed CP and HD.

### Slide 8
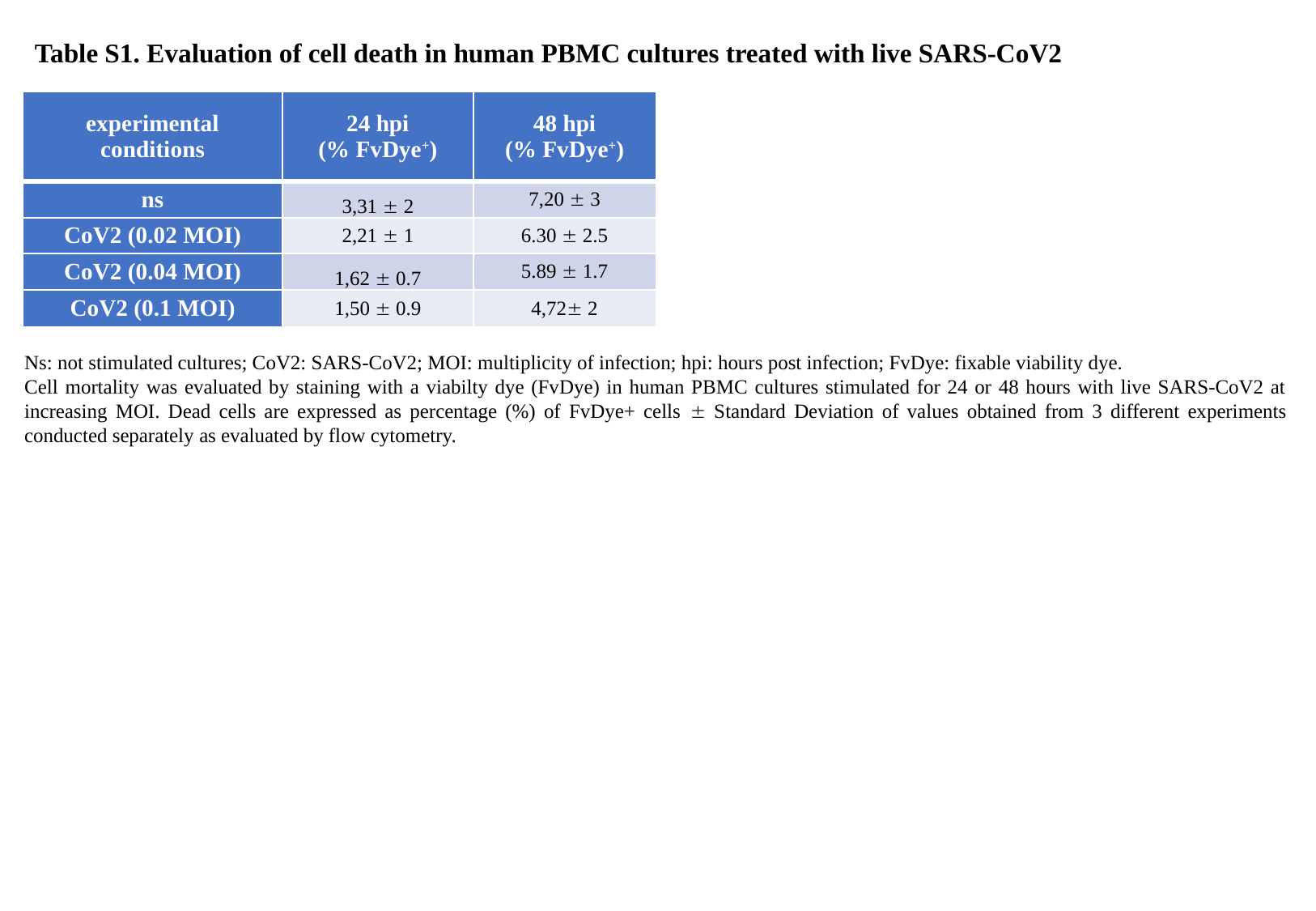

Table S1. Evaluation of cell death in human PBMC cultures treated with live SARS-CoV2
| experimental conditions | 24 hpi (% FvDye^+^) | 48 hpi (% FvDye^+^) |
| --- | --- | --- |
| ns | 3,31 ± 2 | 7,20 ± 3 |
| CoV2 (0.02 MOI) | 2,21 ± 1 | 6.30 ± 2.5 |
| CoV2 (0.04 MOI) | 1,62 ± 0.7 | 5.89 ± 1.7 |
| CoV2 (0.1 MOI) | 1,50 ± 0.9 | 4,72± 2 |
Ns: not stimulated cultures; CoV2: SARS-CoV2; MOI: multiplicity of infection; hpi: hours post infection; FvDye: fixable viability dye.
Cell mortality was evaluated by staining with a viabilty dye (FvDye) in human PBMC cultures stimulated for 24 or 48 hours with live SARS-CoV2 at increasing MOI. Dead cells are expressed as percentage (%) of FvDye+ cells ± Standard Deviation of values obtained from 3 different experiments conducted separately as evaluated by flow cytometry.

### Slide 9
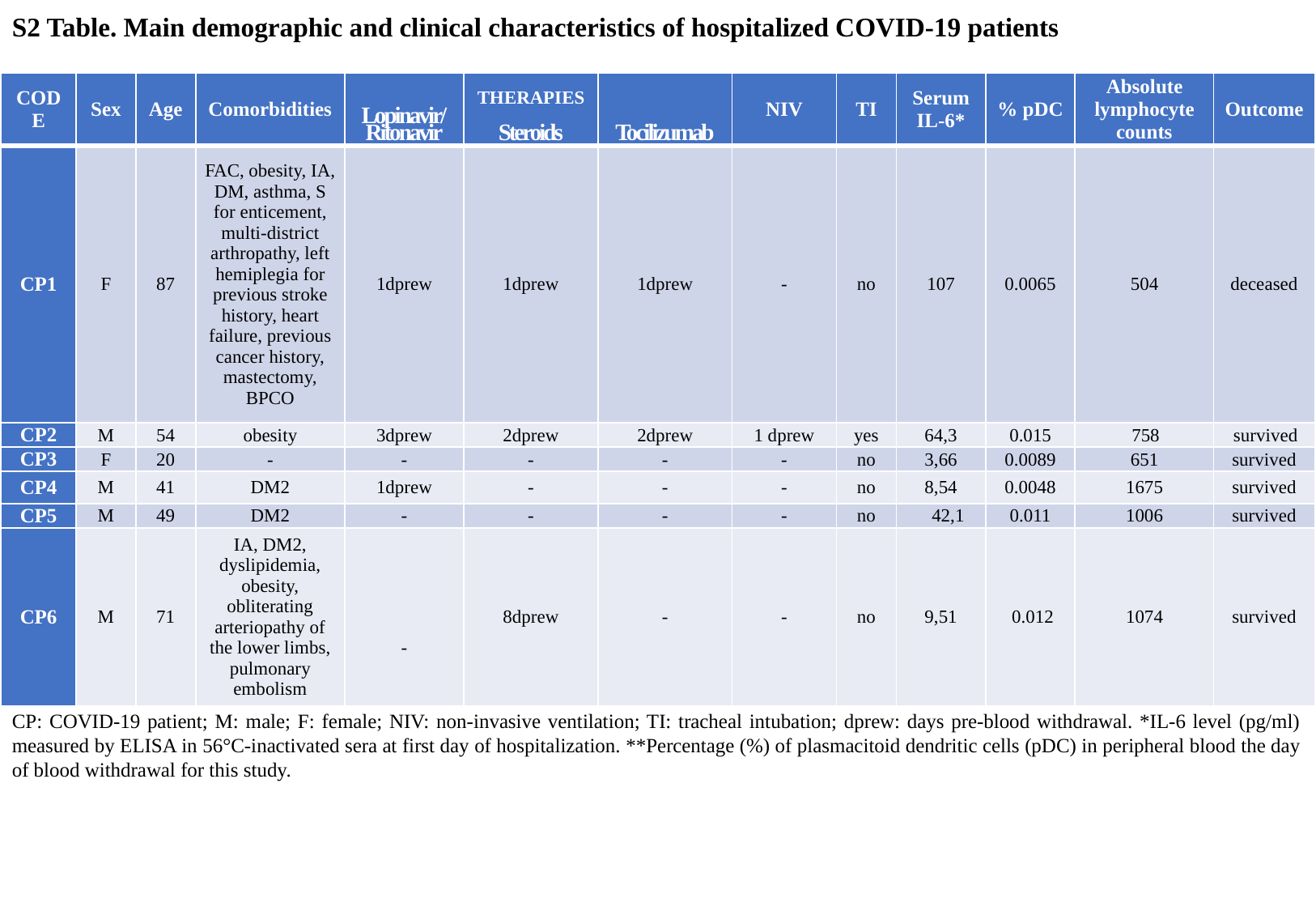

S2 Table. Main demographic and clinical characteristics of hospitalized COVID-19 patients
| CODE | Sex | Age | Comorbidities | Lopinavir/ Ritonavir | THERAPIES Steroids | Tocilizumab | NIV | TI | Serum IL-6\* | % pDC | Absolute lymphocyte counts | Outcome |
| --- | --- | --- | --- | --- | --- | --- | --- | --- | --- | --- | --- | --- |
| CP1 | F | 87 | FAC, obesity, IA, DM, asthma, S for enticement, multi-district arthropathy, left hemiplegia for previous stroke history, heart failure, previous cancer history, mastectomy, BPCO | 1dprew | 1dprew | 1dprew | - | no | 107 | 0.0065 | 504 | deceased |
| CP2 | M | 54 | obesity | 3dprew | 2dprew | 2dprew | 1 dprew | yes | 64,3 | 0.015 | 758 | survived |
| CP3 | F | 20 | - | - | - | - | - | no | 3,66 | 0.0089 | 651 | survived |
| CP4 | M | 41 | DM2 | 1dprew | - | - | - | no | 8,54 | 0.0048 | 1675 | survived |
| CP5 | M | 49 | DM2 | - | - | - | - | no | 42,1 | 0.011 | 1006 | survived |
| CP6 | M | 71 | IA, DM2, dyslipidemia, obesity, obliterating arteriopathy of the lower limbs, pulmonary embolism | - | 8dprew | - | - | no | 9,51 | 0.012 | 1074 | survived |
CP: COVID-19 patient; M: male; F: female; NIV: non-invasive ventilation; TI: tracheal intubation; dprew: days pre-blood withdrawal. *IL-6 level (pg/ml) measured by ELISA in 56°C-inactivated sera at first day of hospitalization. **Percentage (%) of plasmacitoid dendritic cells (pDC) in peripheral blood the day of blood withdrawal for this study.

### Slide 10
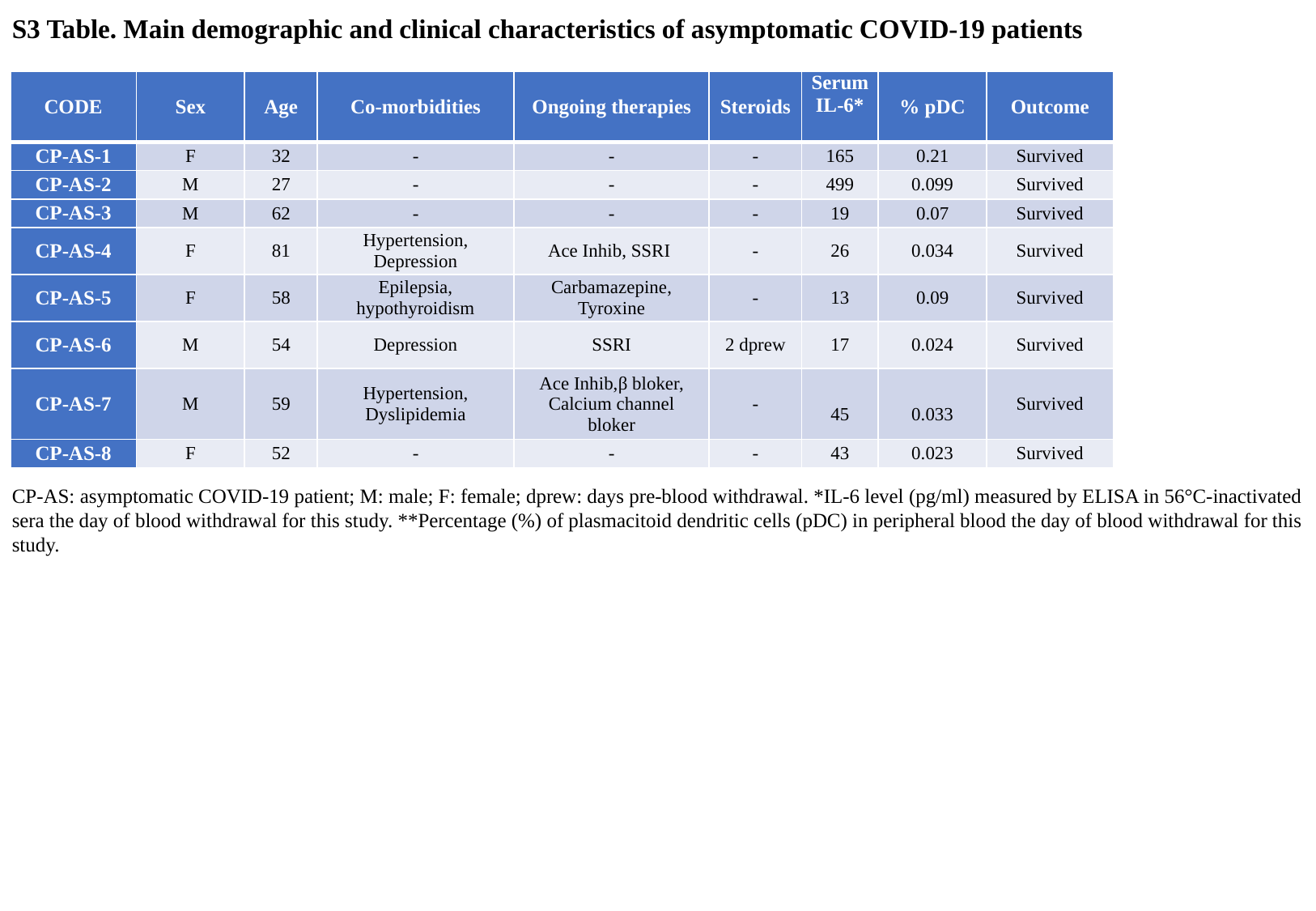

S3 Table. Main demographic and clinical characteristics of asymptomatic COVID-19 patients
| CODE | Sex | Age | Co-morbidities | Ongoing therapies | Steroids | Serum IL-6\* | % pDC | Outcome |
| --- | --- | --- | --- | --- | --- | --- | --- | --- |
| CP-AS-1 | F | 32 | - | - | - | 165 | 0.21 | Survived |
| CP-AS-2 | M | 27 | - | - | - | 499 | 0.099 | Survived |
| CP-AS-3 | M | 62 | - | - | - | 19 | 0.07 | Survived |
| CP-AS-4 | F | 81 | Hypertension, Depression | Ace Inhib, SSRI | - | 26 | 0.034 | Survived |
| CP-AS-5 | F | 58 | Epilepsia, hypothyroidism | Carbamazepine, Tyroxine | - | 13 | 0.09 | Survived |
| CP-AS-6 | M | 54 | Depression | SSRI | 2 dprew | 17 | 0.024 | Survived |
| CP-AS-7 | M | 59 | Hypertension, Dyslipidemia | Ace Inhib,β bloker, Calcium channel bloker | - | 45 | 0.033 | Survived |
| CP-AS-8 | F | 52 | - | - | - | 43 | 0.023 | Survived |
CP-AS: asymptomatic COVID-19 patient; M: male; F: female; dprew: days pre-blood withdrawal. *IL-6 level (pg/ml) measured by ELISA in 56°C-inactivated sera the day of blood withdrawal for this study. **Percentage (%) of plasmacitoid dendritic cells (pDC) in peripheral blood the day of blood withdrawal for this study.
